# Contrasting dates of rainforest fragmentation in Africa inferred from trees with different dispersal abilities

**DOI:** 10.1101/811463

**Authors:** Rosalía Piñeiro, Olivier J. Hardy, Carolina Tovar, Shyam Gopalakrishnan, Filipe Garrett Vieira, M Thomas P Gilbert

## Abstract

The rainforests of Tropical Africa have fluctuated over time. Although today the forest cover is continuous in Central Africa this may have not always been the case, as the scarce fossil record in this region suggests that more arid conditions might have significantly reduced the density of trees during the Ice Ages. Our aim was to investigate whether the dry ice-age periods left a genetic signature on tree species that can be used to date the past fragmentation of the rainforest. We sequenced reduced representation libraries of 182 samples representing five Legume tree species that are widespread in African rainforests and seven outgroups. Phylogenetic analyses identified an early divergent lineage for all species in West Africa (Upper Guinea), and two clades in Central Africa: Lower Guinea-North and Lower Guinea-South. As the structure separating the Northern and Southern clades cannot be explained by geographic barriers, we tested other hypotheses using demographic model testing. The best estimates recovered using ∂a∂I indicate that the two clades split between the Upper Pliocene and the Pleistocene, a date compatible with forest fragmentation driven by ice-age climatic oscillations. Furthermore, we found remarkably older split dates for the shade-tolerant tree species with non-assisted seed dispersal than for light-demanding long-distance wind-dispersed trees. We also show that the genetic diversity significantly declines with the distance from ice-age refugia in the two long-distance dispersed species only. Different recolonisation abilities after recurrent cycles of forest fragmentation seem to explain why we observe congruent genetic spatial structures across species with contrasted timescales.

**SIGNIFICANCE STATEMENT:** Although today the rainforest cover is continuous in Central Africa, the scarce fossil record suggests that arid conditions during the Ice Ages might have reduced the density of trees during the Ice Ages. However, the vast majority of the fossil pollen records preserved in Tropical Africa is too young to inform about this period. Investigating whether the past climate change left a genetic signature on trees can thus be useful to date past forest fragmentation. However, most genetic studies available to date lack resolution as they use limited numbers of loci. In this study we use modern DNA technology to study five Legume trees. Our results show significant differentiation of the populations of each species at a date compatible with forest fragmentation driven by ice-age climatic oscillations. Contrasted timescales were obtained for each species, which probably reflects their different recolonisation abilities after forest fragmentation.

## INTRODUCTION

The rainforest cover in Tropical Africa has fluctuated widely over time. Today the rainforests of West Africa (Upper Guinea) are disconnected from Central Africa (Lower Guinea) by the Dahomey Gap, a forest-savannah corridor along the coast of Benin, Togo and eastern Ghana (Figure 1). However, the fossil record shows that this region was forested under the humid conditions of the last interglacial (from ca. 8,400 years BP) while the current deforestation started only 4,500 years ago following the aridification of the climate (1). It is not clear whether the Dahomey gap also became forested during the previous interglacials, as they do not seem to have been as humid as the last one (2). While the rainforests of Central Africa (Lower Guinea) currently exhibit a continuous distribution, previous genetic studies indicate strong differentiation of tropical trees within the forest. Typically, such structure may be explained by geographic barriers. For instance the river Sanaga, one of the main rivers of Cameroon, runs from inland Cameroon towards the coast delimiting two different subspecies of chimpanzee (3) and also creating a deep genetic divergence of mandrill populations on both sides (4). In Gabon, the Oougué river acts as an effective barrier for dispersal of mandrills (4) and gorillas(5). In contrast to primates the genetic structure of tropical trees cannot be explained by effective barriers to dispersal such as the main rivers and mountain chains in this area (6, 7).

**Figure 1.**
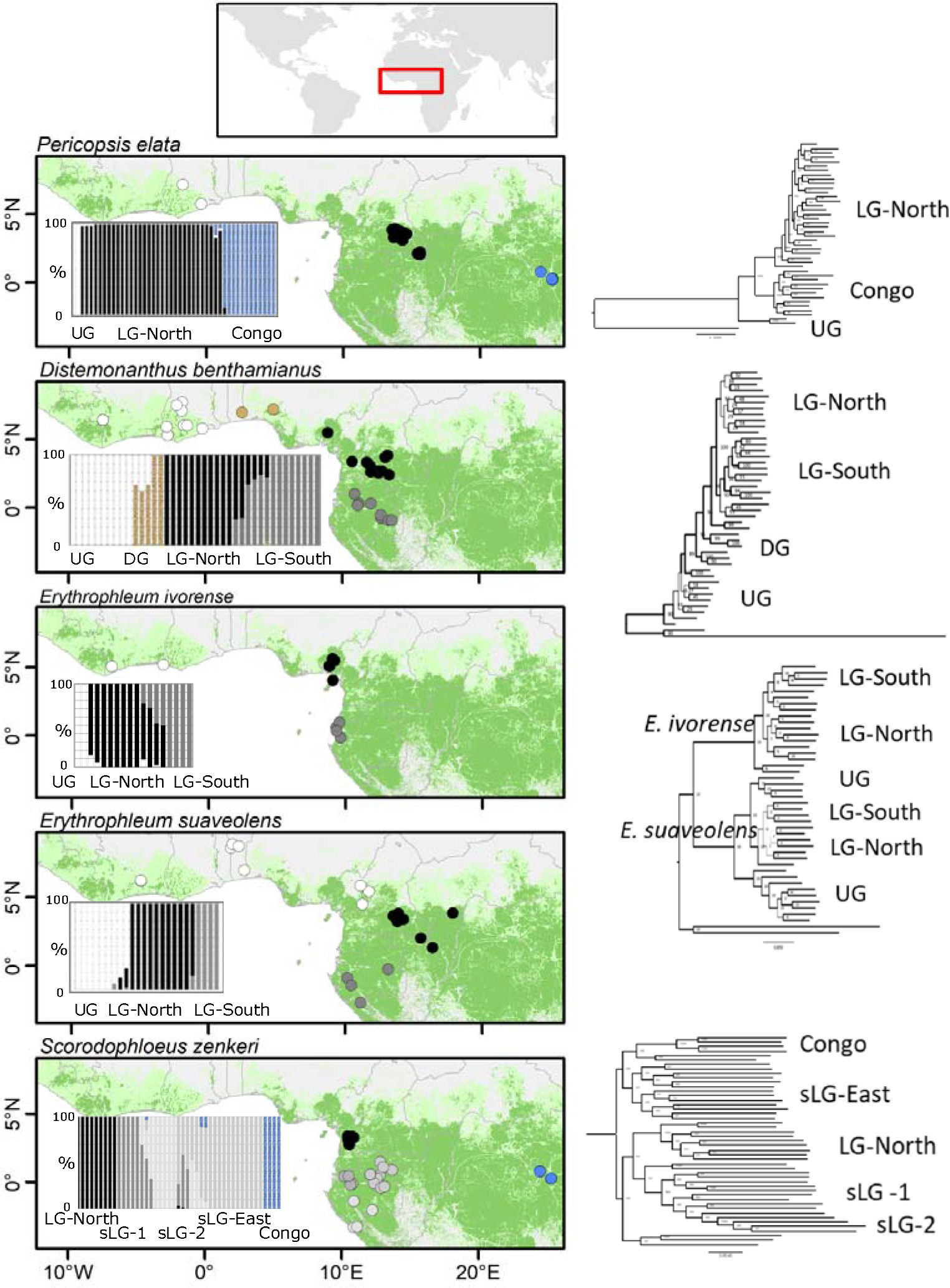
Genetic ancestry and phylogenetic reconstructions for *Pericopsis elata, Distemonanthus benthamianus, Erythrophleum ivorense, E. suaveolens*, and *Scorodophloeus zenkeri* based on Genotyping by Sequencing (GBS). Left: ADMIXTURE analyses. Barplots show the probability of assignment of individuals to genetic clusters. Centre: geographic distribution of individual trees according to the genetic cluster they belong to (admixed individuals with <70% assignment probability to a single genetic cluster have been removed). Right: RAxML phylogenetic analyses using the GBS genotype calls with outgroups. Branch width is proportional to bootstrap supports. Green areas in map represent present-day rainforest cover (54) (see also S3).

Recent studies based on chloroplast DNA, nuclear microsatellites, and low-copy nuclear genes (6–8) suggest that the observed historical isolation of the tree populations in Tropical Africa was caused by forest fragmentation during the cold and dry Ice-Age periods, which occurred on several cycles, along the Pleistocene (9). However, dating the fragmentation of the rainforest and determining where the ancestral populations of each species was, has been challenging due to the low number of markers investigated. For example with regards to dating, three prior studies have attempted to estimate the divergence of tree populations in the Pleistocene, although with large uncertainty due to the low numbers of molecular markers used (10–12). A fourth study on the genus *Greenwayodendron* places the divergence estimates in the Pliocene/Pleistocene(13). As for the location of forest fragments that allowed tropical tree species to survive the Ice Ages, these have been postulated based on the fossil record and palaeoclimatic reconstructions. Areas that harbour high species richness have been proposed as refugia (Figure 2), assuming declines in the number of species outside the hypothesized refugia (14). Likewise, declines of genetic diversity with distance from refugia are expected to result from recolonisation after forest fragmentation. Despite the potential of genetic diversity gradients to help locate the areas in which forest species survived, the available genetic data lack resolution to assess diversity gradients over space.

**Figure 2.**
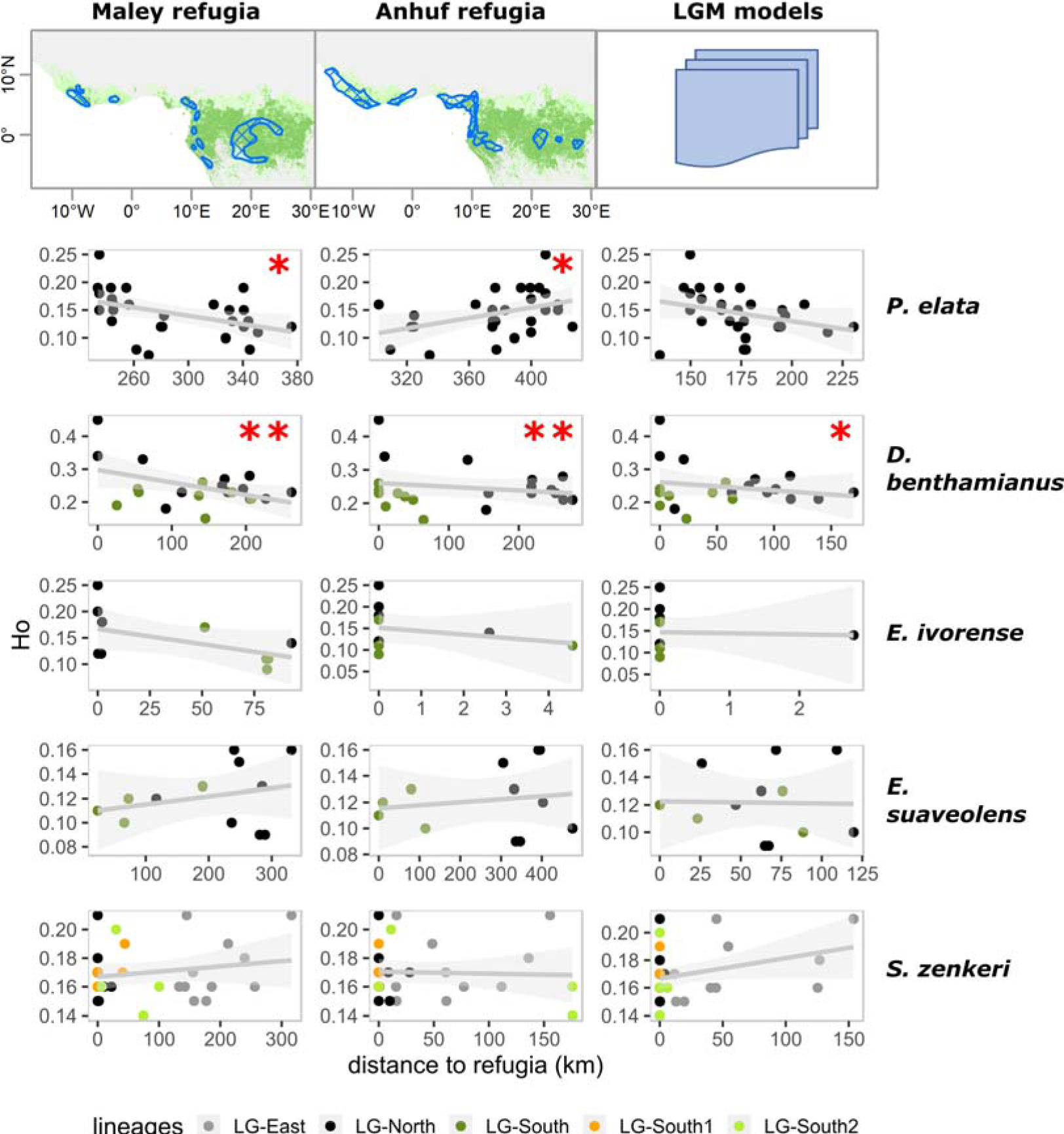
Decline of genetic diversity, based on Genotyping by Sequencing, with distance from refugia in Central Africa for *Pericopsis elata, Distemonanthus benthamianus, Erythrophleum ivorense, E. suaveolens*, and *Scorodophloeus zenkeri*. Linear regression of the genetic diversity -observed heterozygosity, H_o_- of each genotyped individual tree against the geographical distance from the closest Last Glacial Maxima refugia ca. 20,000 yr. BP, and the genetic cluster as a random variable in order to account for genetic diversity differences across genetic lineages. Admixed individuals with less than 70% genetic ancestry to a single group were excluded. Three different hypotheses of LGM forest refugia were considered. The forest refugia postulated by Maley and Anhuf, based on palaeoclimatic and palynological data: (i) LGM-Maley (14), and (ii) LGM-Anhuf, (27). In addition, specific forest refugia based on the LGM niche models of each species were tested (S7): (iii) LGM-species niche models (see methods).

To overcome these challenges, we generated reduced representation genomic data using Illumina sequencing technology, for five Legume rainforest tree species, in order to investigate the genetic signal of changes in the rainforest cover during the Ice Ages. All five species -*Pericopsis elata* (Harms) Meeuwen, *Distemonanthus benthamianus* Baill. *Erythrophleum ivorense* A. Chev., *Erythrophleum suaveolens* (Guill. & Perr.) Brenan, and *Scorodophloeus zenkeri* Harms- are widespread in the African rainforest (Figure 1) and exhibit differences in light tolerance and dispersal capacity (S1). Based on their ecology and dispersal biology, *P. elata* and *D. benthamianus* are the most adapted to long-distance colonization. They are both light-demanding and wind-dispersed, with significant long-distance dispersal events (15). The two *Erythrophleum* species are light-demanding and besides its primary ballistic dispersal they exhibit secondary dispersal of their seeds by animals, but no evidence of long-distance dispersal has been detected in direct measurements with molecular markers (15). *S. zenkeri*, a strict shade-tolerant species with seed dispersal non-assisted by wind or animals, exhibits the most limited colonising capacity.

In particular we aimed to solve the following questions:

- Are the tree populations in West African rainforests (Upper Guinea) well-differentiated from Central African rainforests (Lower-Guinea) or did the expansion of the rainforest during the humid Holocene favoured dispersal and geneflow between the two forest blocks?
- Are the divergence times estimated by sequencing large portions of the genome compatible with isolation and fragmentation of the Central African rainforest during the dry and cold Ice Ages?
- Can we trace the genetic signal of recolonisation of tree populations from putative glacial refugia?
- What are the similarities and differences in the patterns of genetic diversity and differentiation in the light of different ecologies and dispersal capacities of the five tree species?

## RESULTS

A total of 182 samples were successfully sequenced (175 samples of the five study species and 7 outgroup taxa). Samples received an average number of reads between 3,387,236 in the *P. elata* and 5,518,179 in *D. benthaminanus* (S2). Our protocol compensated the limitation of uneven coverage across loci typical of GBS studies by sequencing two libraries per individual.

For the GBS genotype calls without outgroups done with TASSEL between 10,665 (*D. benthamianus*) and 27,838 SNPs (*E. suaveolens*) were present in at least 50% of individuals. For the genotype likelihood framework we retained between 568,240 reads in *P. elata* (16 depth coverage) and 1,561,119 reads in *D benthamianus* (32X coverage). The number of SNPs ranged from 17,921 in *D. benthamianus* to 70,295 in *S. zenkeri* for the less stringent quality cut-off filtering 1 (Table 1, S2).

**Table 1:** Demographic history of *Distemonanthus benthamianus, Erythrophleum ivorense, E. suaveolens*,and *Scorodophloeus zenkeri* using ∂a∂I.

|  | Dataset FILTERING 1 |  |  |  | Dataset FILTERING 2 |  |  |  |
| --- | --- | --- | --- | --- | --- | --- | --- | --- |
|  | *D. benthamianus* | *E. ivorense* | *E. suaveolens* | *S. zenkeri* | *D. benthamianus* | *E. ivorense* | *E. suaveolens* | *S. zenkeri* |
| **Divergence time (Kyr. ago)** |  |  |  |  |  |  |  |  |
| nLG – sLG | 28.98 | 103.06 | 359.28 | 3436.99 | 7.70 | 76.90 | 101.78 | 3085.94 |
| UG – (nLG, sLG) | 1224.79 | 1151.31 | 3493.44 |  | 115.53 | 1034.19 | 1053.05 |  |
| sLG1, sLG2 |  |  |  | 1462.52 |  |  |  | 174.03 |
| **Model details** |  |  |  |  |  |  |  |  |
| Log-likelihood | -15764.60 | -394.29 | -25466.29 | -17122.81 | -11098.82 | -637.26 | -14168.11 | -13590.11 |
| # of SNPs | 17921.38 | 29513.13 | 61667.59 | 70271.16 | 11323.50 | 22151.64 | 35042.61 | 54156.53 |

Divergence time in 1000 years are shown for the best-fitting model under Isolation with Migration. Based on the admixture and phylogeny results, for the three species widespread in Upper and Lower Guinea -*D. benthamianus*, *E. suaveolens*, *E. ivorense*-, we fitted a three-population model with the tree topology (Upper Guinea-UG (Northern Lower Guinea-nLG, Southern Lower Guinea-sLG). In addition to a model with symmetric migration among the three populations, we fitted a model with no migration between UG and sLG after the split. Note that the best fitting-model for the three species does not include a migration component between UG and sLG. For *S. zenkeri*, absent in Upper Guinea, we estimated split times based on the tree topology (Northern Lower Guinea-nLG, (Southern Lower Guinea 1-sLG1 and Southern Lower Guinea2-sLG2)). The demographic parameter estimates are provided for two datasets per species, depending on the stringency of the filtering criteria, filtering 1, the most lenient and filtering 2, the intermediate filtering. Filtering 3 is not shown, since it resulted in far fewer variable sites than filtering 1 and 2 (see full results in S5).

### Genetic Ancestry and Phylogenetic reconstructions

#### Upper Guinea and Lower Guinea

We conducted RAxML phylogenetic analyses for each species using the GBS genotype calls with outgroups (Figure 1, Right; S3). For the species present in West Africa (*P. elata, D. benthamianus, E. ivorense*, and *E. suaveolens*) the Upper Guinean populations clustered in independent clades that were sister to the Lower Guinean clades. Similarly, ADMIXTURE barplots of probability of assignment of individuals to populations (Figure 1, Left) showed separate Upper Guinean (UG) genetic clusters when the number of ancestral populations (K) was set to K=3 (*E. ivorense, E. suaveolens*, and *P. elata*) or K=4 (*D. benthamianus*). In the case of the species widespread in Lower Guinea (LG) -*D. benthamianus, E. ivorense, E. suaveolens*, and *S. zenkeri*- two reciprocally monophyletic clades stand out for LG-North and LG-South (Figure 1, Right; S3). Similarly, in the ADMIXTURE analyses a split between the LG-North and the LG-South genetic clusters was observed (Figure 1) at K=3 for *E. ivorense* and *E. suaveolens*, and at K=4 for *D. benthamianus* and *S. zenkeri*. The two species sampled in Congo (P. elata, and S. zenkeri) revealed independent genetic groups based on RAxML and ADMIXTURE analyses. However, the geographic coverage of our samples in this biogeographic region is incomplete.

#### D. benthamianus

The ADMIXTURE analysis at K=2, where the minimum cross validation (CV) was found, revealed an UG and a LG cluster, with genetically intermediate samples in the Dahomey Gap. At K=3 the DG is retrieved as an independent cluster, with admixture from the UG region in the West. At K=4 in addition to the UG, and DG clusters, LG splits into two clusters LG-North and LG-South, with ancestry shared between the two groups in the geographically intermediate areas. From K=5 the samples with shared ancestry between LG-North and LG-South form an independent cluster. From K=>6 further genetic subgroups are found within the UG and LG-North, with no geographical congruence. The rooted ML phylogenetic tree without admixed individuals agrees with K=4. It consists of several basal clades for UG, two reciprocally monophyletic well-supported clades in LG–North and LG-South, and an intermediate clade for the DG. The admixed individuals fell in between the main clades.

#### E. ivorense

The ADMIXTURE analyses do not show a clear minimum in CV values. At K=2 samples from UG and LG-North clustered together in one group and LG-South samples in a second cluster, with admixed individuals between the LG-North and the LG-South. At K=3 the clustering corresponds to UG, LG-North, and LG-South, with admixed individuals in between the two latter. From K=4 and higher, genetic subgroups were distinguished within the LG-North and the LG-South clusters, with no coherence across K. The rooted RAxML phylogeny revealed three main clades in agreement with K=3: a basal clade in UG, and two sister clades corresponding to LG-North and LG-South. The admixed individuals between LG-North and LG-South were placed together with the Southern cluster.

#### E. suaveolens

The ADMIXTURE analyses split an UG and a LG clusters at K=2, where the minimum CV is reached. At K=3 the genetic clusters retrieved were UG, LG-North, and LG-South. From K=4 and higher, subgroups appear randomly within UG, and from K=6 and higher within LG-North. The RAxML phylogenetic tree retrieves three clades corresponding to UG, LG-North and LG-South. No individuals with shared ancestry between groups were detected. In this species the UG cluster does not correspond to the West African Guineo-Congolian forests but to gallery forests embedded in West African savannahs, including the forest-savanna mosaic of Cameroon, XI. Guinea-Congolia/ Sudania regional transition zone (16).

#### S. zenkeri

No clear minimum CV was shown in the ADMIXTURE analyses. At K=2 LG-South splits from the rest of the samples. At K=3 the following clusters are observed: LG-North, LG-South, and a LG-east/Congo, with significant admixture between LG-South and LG-east/Congo. At K=4 the LG-east and the Congo clusters are retrieved as independent groups. At K=5 the LG-South is split into a Northern (LG-South1) and a Southern cluster (LG-South2), with admixed individuals between them, and also with LG-East. From K=6 and higher random genetic groups arise within LG-South1, LG-South2 and LG-East.

#### P. elata

At K=2 LG-North splits from UG/Congo (minimum CV). At K=3 UG, LG-North, and Congo were revealed. From K=4 random subgroups within LG-North, and Congo, are retrieved. The rooted RAxML phylogenetic tree revealed three well-supported clades in agreement with K=3, where the UG clade is basal with respect to the two LG sister clades.

### Demographic inference

We inferred the demographic history of each species using *∂a∂I* (Table 1, S4, S5). The fit substantially improved when considering a scenario of no migration between Upper Guinea and South Lower Guinea after divergence, in all cases except for one of the *E. ivorense* scenarios. For each scenario, the two different levels of data filtering gave different estimates but congruent for each species. The scenarios with more SNPs (filtering level 1) gave older split dates while the scenarios with fewer SNPs (filtering level 2) fitted slightly more recent splits. The most restrictive filtering (filtering level 3) yielded fewer than 10,000 SNPs in all cases so estimates are not reliable.

For the divergence between LG-North and LG-South, *∂a∂i* produced different estimates across species (Table 1). The most likely models estimated the most recent split ca. 29 Kyr. BP for *D. benthamianus*, followed by *E. ivorense* ca. 103 Kyr BP and *E. suaveolens* ca. 359 Kyr. BP. The oldest was in *S. zenkeri* ca. 3.5 Myr. BP. The scenarios built using filtering level 2 generally inferred younger North-South splits but the differences across species persisted (ca. 7.6 Kyr in *D. benthamianus*, ca. 77 Kyr. BP in *E. ivorense*, ca. 102 Kyr. BP in *E. suaveolens*, and ca. 3.1 Myr BP in *S. zenkeri*).

### Gradients of genetic diversity over space

We traced the spatial signal of recolonisation after forest fragmentation, by relating genetic diversity of individuals per species with distance from the three hypothesised LGM forest refugia, accounting for differences across gene pools (Figure 2). We found a significant negative relationship between observed heterozygosity, H_o_, and distance to refugia only for *D. benthamianus* and *P. elata* (Figure 2, Table 2). Higher genetic diversity is found in individuals of *D. benthamianus* that are closer to the LGM forest refugia: LGM-Maley (p<0.01), LGM-Ahnuf (p<0.01), and LGM-species niche model (p<0.05). The present-day distribution of *D. benthamianus* is mostly closer to the coast rather than in the core of the Congo forest (Figure 1), All three postulated refugia suggest large areas of forest survival during the LGM along the coasts of Cameroon and Gabon thus, the observed gradient in H_o_ is dominated by the distance to the Coastal refugia rather than to the Congo refugia (Figure 2). For *P. elata* we detected a significant decline of the genetic diversity as individuals are located further away from LGM-Maley (p<0.05). As the distribution of *P. elata* lays between the Coastal and the Congo refugia (Figure 1, S6), we tested the relationship between H_o_ and the distances to both refugia separately. Here we found a significant negative relationship with distance to the LGM-Maley-Congo (p<0.01) and a positive relationship with scenarios LGM-Maley-Coastal. We also found a positive significant relationship with LGM-Anhuf (Figure 2, Table 2), where coastal refugia is closer to current *P. elata* populations. Finally, the LGM-species niche model, which shows only inland LGM refugia restricted to NE Cameroon, does not exhibit any significant relationship.

**Table 2:**
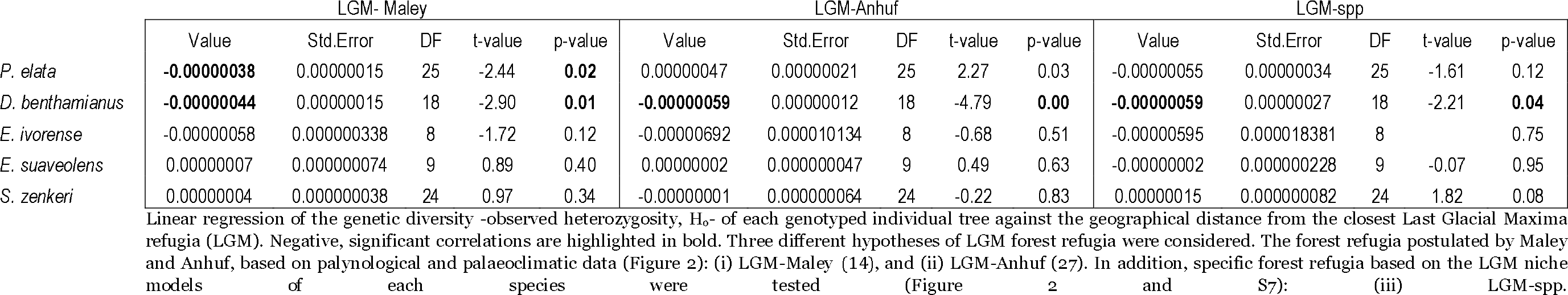
Decline of genetic diversity with distance from refugia in Central Africa (Lower Guinea) for *Pericopsis elata, Distemonanthus nthamianus, Erythrophleum ivorense, E. suaveolens*, and *Scorodophloeus zenkeri*.

|  | LGM- Maley |  |  |  |  | LGM-Anhuf |  |  |  |  | LGM-spp |  |  |  |  |
| --- | --- | --- | --- | --- | --- | --- | --- | --- | --- | --- | --- | --- | --- | --- | --- |
|  | Value | Std.Error | DF | t-value | p-value | Value | Std.Error | DF | t-value | p-value | Value | Std.Error | DF | t-value | p-value |
| *P. elata* | **-0.00000038** | 0.00000015 | 25 | -2.44 | **0.02** | 0.00000047 | 0.00000021 | 25 | 2.27 | 0.03 | -0.00000055 | 0.00000034 | 25 | -1.61 | 0.12 |
| *D. benthamianus* | **-0.00000044** | 0.00000015 | 18 | -2.90 | **0.01** | **-0.00000059** | 0.00000012 | 18 | -4.79 | **0.00** | **-0.00000059** | 0.00000027 | 18 | -2.21 | **0.04** |
| *E. ivorense* | -0.00000058 | 0.000000338 | 8 | -1.72 | 0.12 | -0.00000692 | 0.000010134 | 8 | -0.68 | 0.51 | -0.00000595 | 0.000018381 | 8 |  | 0.75 |
| *E. suaveolens* | 0.00000007 | 0.000000074 | 9 | 0.89 | 0.40 | 0.00000002 | 0.000000047 | 9 | 0.49 | 0.63 | -0.00000002 | 0.000000228 | 9 | -0.07 | 0.95 |
| *S. zenkeri* | 0.00000004 | 0.000000038 | 24 | 0.97 | 0.34 | -0.00000001 | 0.000000064 | 24 | -0.22 | 0.83 | 0.00000015 | 0.000000082 | 24 | 1.82 | 0.08 |

In the remaining species *E. ivorense, E. suaveolens*, and *S. zenkeri* no significant relationships were found (Figure 2, Table 2, S6). In the case of *E. ivorense* the results need to be taken with caution as low sampling sizes may have reduced statistical power.

## DISCUSSION

For the five rainforest tree species investigated, at least three main intraspecific lineages were identified in the phylogenetic analyses. An early divergent lineage in West Africa (Upper Guinea, UG) was detected in all species occurring in this forest block (*D. benthamianus, E. suaveolens, E. ivorense*, and *P. elata*). For all species widespread in Central Africa (Lower Guinea, LG) two lineages were retrieved: Northern Lower Guinea, LG-north) and a Southern Lower Guinea, LG-south (*D. benthamianus, E. suaveolens, E. ivorense*, and *S. zenkeri*).

### Early divergence in West African rainforests

The finding of early divergent lineages in West Africa for all the species is in tune with the hypothesis that Upper Guinea is an independent biogeographic region with numerous endemic species (8, 16–18). Although the fossil record indicates that the Dahomey gap was forested in the last interglacial, and thus Upper and Lower Guinea were connected between 8,400 and 4,500 years ago (1, 19), our data show that no genetic homogenisation between the two forest blocks occurred. It is likely that previous interglacials were less humid than the last one so there is no guarantee that the Dahomey Gap became forested during those periods (2). This seems to have favoured the long-term differentiation of the two forest blocks. To our knowledge this is the first study that estimates well-supported intraspecific phylogenies between Upper and Lower Guinean plant populations using outgroups in order to estimate the direction of evolution. Based on relatively small numbers of markers, early divergent lineages, c.a. 500 Kyr. BP, have also been found in west African populations of chimpanzees (3, 20, 21) and woodpeckers (22). Studies on other African lowland rainforest birds (23–25), forest-dwelling rodents (26) and African bushbucks suggest that haplotypes are rarely shared between populations sampled across the Dahomey Gap although the relationships between clades is not well resolved.

### North-South Genetic differentiation in Central Africa suggests Rainforest Fragmentation during the Ice-Ages

For each species, our phylogenetic analyses identified northern and southern lineages in Central Africa that do not correspond to any geographic barrier or current discontinuity in the distribution of the forest. These results are consistent with the genetic structuring of other tropical trees in Lower Guinea (6, 7), as well as with the ADMIXTURE analyses, which provided additional insight into the levels of admixture among the central African lineages of each species. While *D. benthamianus* and *E. ivorense* showed admixture between the North and the South, *E. suaveolens* and *S. zenkeri* exhibited sharp differentiation and no admixed individuals between the two regions.

Using *δαδl* to reconstruct the population history of the four species present in North and South Lower Guinea, our best-fit comparison consistently supported models involving North-South divergence and subsequent gene flow. We also noticed that the fit substantially improved when considering no migration between Upper Guinea and South Lower Guinea after divergence. These clades diverged within the Upper Pliocene and the Pleistocene, with the oldest genetic split found in *S. zenkeri* ca. 3.4-3.1 Myr. BP, and the most recent in *D. benthamianus* ca. 29,000-7,7oo yr. BP. Our time estimates indicate North-South differentiation of the forest populations during the dry glacial climatic periods that took place from the Pliocene and especially during the Pleistocene (9). Altogether, our data are compatible with the differentiation of the genetic lineages in Northern and Southern Lower Guinea as a result of forest fragmentation during the dry glacial periods and subsequent admixture as a result of forest expansion during the humid interglacial periods.

#### Decline of genomic diversity from putative glacial refugia

We traced the spatial signal of recolonisation during the humid periods, by examining genetic diversity gradients over space. We found significant declines of genetic diversity from coastal refugia in *D. benthamianus* and from inland Congo refugia in *Pericopsis elata*. This finding suggests the survival of *D. benthamianus* in coastal refugia in Cameroon and Gabon, although the exact location cannot be determined since no differences were revealed among the different hypotheses of LGM forest refugia considered based on the palaeoclimatic reconstructions (14, 27) or estimated from the potential climatic distribution of each species during the LGM. In the case of *P. elata* survival in inland forest refugia postulated by Maley in the Congo Basin is highly probable. For *S. zenkeri* and *E. suaveolens* no significant declines of the genetic diversity were found.

#### Genomic diversity and differentiation and dispersal capacities

Our estimates of admixture and split dates are consistent with prior knowledge of the biology of the study species. In particular, with two traits that may be key for colonising new areas after forest fragmentation: light tolerance and dispersal capacity. Long-distance dispersal of the seeds is especially relevant in the case of early successional communities and expanding populations, as it not only transports seeds over very long distances but also generates establishment opportunities. *∂a∂i* detected more recent splits between the North and South in *D. benthamianus*, than for the two *Erythrophleum* species. The oldest split was detected in the shade-tolerant, non-assisted dispersed species *S. zenkeri*. The fact that we detected more recent signals of North-South fragmentation in the species with long-distance dispersal capacity suggests that the old signals of fragmentation may be more easily erased in these species than in species with limited dispersal like *S. zenkeri*. The ability of long-distance dispersal may have made a difference over subsequent cycles of forest fragmentation and recolonisation that took place during the Pleistocene. Our results also show a decline of genetic diversity from forest refugia in long-distance-dispersed species only. While *D. benthamianus* likely survived in coastal refugia, *P. elata* probably did so in inland refugia in the Congo Basin. The fact that we were able to trace the genetic significant declines of genetic diversity outside refugia in the long-distance dispersed species suggests that we may be detecting the signal of a recent dispersal for those species.

### Conclusions

GBS data of five Legume tree species widespread in African rainforests reveal: i) early divergence of the West African populations (Upper Guinea) from Central Africa (Lower Guinea), and ii) a clear North-South differentiation in Lower Guinea despite the absence of discontinuities in the rainforest cover or other geographic barriers, such as rivers and mountain chains. However, divergence times vary widely among species, from the Pliocene for shade-tolerant trees with non-assisted seed dispersal to late Pleistocene or Holocene for pioneer long-distance wind-dispersed trees. We conclude that different responses of tree species to recurrent forest fragmentation cycles driven by past climate fluctuations may explain why we observe congruent genetic spatial structures with contrasted timescales. Species with higher colonising abilities seem to have been able to erase old signals of genetic differentiation compared to species with limited dispersal and light tolerance.

## MATERIALS AND METHODS

### Study species: dispersal capacity, light tolerance, reproductive system

*Pericopsis elata* is a light‐demanding, pioneer or non-pioneer and wind-dispersed. Seeds disperse on average 214 m with a very flat tail. Most seeds disperse less than 100m, but a significant amount of seeds experience long distance dispersal (>1 km) (Olivier Hardy, personal observation). *D. benthamianus* is a pioneer light-demanding and wind-dispersed species (28). Individual trees are not aggregated in the field. It is an indicator of disturbed rainforests. Although most of the *D. benthamianus* seeds disperse over short distances (c. 70 m), 30% of seed immigration was detected (15). This indicates that the distribution of dispersal distances is very flat tailed, indicative of long-distance dispersal events. The two *Erythrophleum* species are light-demanding (28), the coastal congener *E. ivorense* is a pioneer while *E. suaveolens* is a non-pioneer (Anais Gorel, personal communication). The fruits exhibit primary ballistic dispersal and secondary dispersal by primates, *Cephalophus* species and rodents has been reported (29). The distribution is non gregarious, and they are indicators of secondary rainforests. Based on genetic markers seed dispersal distances of 210 m were detected in *E. suaveolens*, but long distance dispersal events seem rare (30). *Scorodophloeus zenkeri* is a shade tolerant species that exhibits ballistic dispersal through the explosion of the pods (R Piñeiro 2017, personal observation). Trees are locally aggregated in the field. It requires high environmental humidity (31) and is an indicator of undisturbed rainforests.

### DNA extraction and Genotyping by Sequencing

Leaf and cambium material were collected in the rainforests of West and Central Africa between 2005 and 2014 (S8). The samples were immediately dried with silica-gel in order to preserve the DNA quality. Between 18 and 46 samples of each species were selected in order to represent their distribution in Central Africa, with special emphasis in the biogeographic region of Lower Guinea. A few samples from the less accessible rainforest of the Congo Basin were included. Outgroup taxa (S8) were selected based on available legume phylogenies (32, 33).

Overall 362 GBS libraries, from 182 individuals, were initially sequenced on four Illumina lanes (HiSeq2000 San Diego, CA, USA), using 100-bp Single Read chemistry. For each library two DNA extractions were performed using the DNeasy Plant Minikit columns (Qiagen), and pooled in order to generate sufficient DNA for the GBS protocol. One blank per plate was included. DNA quality was checked on a 1.5% agarose gel and DNA quantity was measured with Qbit HS (Life technologies, Grand Island, NY). The DNA was purified with a ZR-96 DNA Clean up kit (Zymo Research Corp.). Subsequently, genotyping by Sequencing (GBS) was performed at the Genomic Diversity and Computational Biology Service Unit at Cornell University (Ithaca, NY) according to a published protocol (34). One microgram of DNA of each species was initially used in order to optimise the GBS protocol, in particular aiding the choice of the most appropriate restriction enzyme. Specifically, three libraries were built for each species using three different enzymes: ApeKI (4.5-base cutter), EcoT22I and PstI (both 6-base cutters) and checked for appropriate fragment sizes (<500bp) and distribution on an Experion automatic electrophoresis system (Bio-Rad laboratories, USA). Given these results, we elected to use the enzyme EcoT22I for subsequent data generation, as it yielded appropriate fragment sizes (<500bp) and distributions for all study species.

### Genotype Calling and Site Frequency Spectrum

We limited the impact of uneven coverage of samples typical for GBS data by building and sequencing two independent libraries for each individual. Two complementary bioinformatic pipelines were implemented for genotype calling and estimating summary statistics used for downstream analyses.

#### Genotype calls without outgroups

For those analyses that require single nucleotide polymorphism (SNP) and genotype calls, we used the Universal Network-Enabled Analysis Kit (UNEAK) pipeline (35) within the software TASSEL 3.0 (36), suitable for analysis of GBS data at the intraspecific level in the absence of a reference genome. Reads were trimmed to 64 bps (to avoid sequencing errors at the ends of reads) and identical reads collapsed into tags (i.e. alleles). Tag pairs having a single base pair mismatch were identified as candidate loci and used for SNP calling. Tag pairs forming complicated networks, likely to result from repeats, paralogs and sequencing error, were filtered out. In order to filter out false-positive SNPs: (i) the minimum number of reads per GBS tag (i.e. allele) was set to five (ii), an error tolerance rate (ETR) of 0.05 was established, and (iii) SNPs with a genotype missing rate >50% were removed.

#### Genotype likelihood framework for analyses with outgroups

Building a “reference” catalog: As the first step in estimating genotype likelihoods at variable sites across the sequenced loci, we constructed a reference catalog against which we could align the GBS reads. We used the tags built with TASSEL to obtain 64 bp long reads. Subsequently, the query read sequences from TASSEL were concatenated with a spacer of 200 Ns to obtain the reference.

##### Genotype likelihood computation

For many of the downstream analyses, a genotype calling approach from NGS data may bias the population genetic estimates (37). Therefore we compute the genotype likelihoods at all the variable sites using the software Analysis of Next-Generation Sequencing Data, ANGSD v0.914 (38). First, we mapped the GBS reads from each sample to the constructed reference catalog using the PALEOMIX pipeline (39). As part of this pipeline, AdapterRemoval v2 (40) was used to trim adapter sequences, merge the paired reads and discard reads shorter than 30 bp. BWA v0.7.15 was then used to map the processed reads to the reference (41). Minimum mapping quality was set at 15 while minimum base quality was set to five. Subsequently, genotype likelihoods were calculated in ANGSD using the following filters, (i) the baq option was used to recompute base alignment qualities, allowing us to reduce false SNPs due to misalignment, (ii) a Hardy-Weinberg Equilibrium (HWE) p-value greater than 0.001, and (iii) the total coverage at any site cannot exceed 55 times the number of samples. The depth distribution was plotted for each species and a cut-off was chosen in order to exclude outlier sites at extremely high coverage. Given the diversity in our reads, the ‒C filter, downgrading mapping quality for reads containing excessive mismatches, was not used.

##### Site Frequency Spectrum

Multi-population Site Frequency Spectrum (SFS) was estimated from the genotype likelihoods using the utility programme realSFS provided with ANGSD. Three different quality cut-offs were used to compute three different SFS for each dataset: (filtering 1) minimum mapping and base quality were both set to 15, and only sites where at least 50% of the samples were covered by one or more reads were retained; (filtering 2) minimum mapping and base quality were both set to 10, and only sites where at least 65% of the samples were covered by one or more reads were retained; (filtering 3) minimum mapping and base quality were both set to 30, and only sites where at least 75% of the samples were covered by one or more reads were retained.

### Inference of genetic clusters

The software Admixture v.13 was used to calculate the probability of assignment of individuals to genetic clusters (42) from the genotype calls datasets without outgroups. Individuals exhibiting admixture between clusters were detected. For each species, K = 1 to 12 genetic clusters were tested with 20 randomly seeded replicate runs at each K and five-fold cross-validation (CV). Barplots showing the probability of assignment of individuals to the genetic clusters were generated in R using the package ggplot2.

The number of K that best fits the data was estimated using the lowest five-fold CV. Since these values did not always work appropriately due to low sample sizes in some of the clusters (43), we used (i) the congruence of the genetic clusters with the geography, and (ii) the congruence of the genetic clusters with the phylogenies as additional criteria to select the optimal K.

### Phylogenetic relationships

Maximum likelihood phylogenies were built for each species using RAxML 8.0 (44) from the Genotype Calls datasets with outgroups. Rooting with outgroups allowed us to assess the direction of evolution. All SNPs were concatenated into a single alignment, with missing data (Ns) and heterozygous positions entered as needed. Bootstrap support was calculated from 100 replicate searches with random starting trees using the GTR+ gamma nucleotide substitution model.

### Demographic inference with δαδl

Population demographic models were estimated using δαδl (45), a method that uses the SFS of populations to infer their demographic history. Based on the admixture and phylogeny results, for the three species widespread in Upper and Lower Guinea -*D. bentamianus, E. suaveolens, E. ivorense*-, we fitted a three-population model with the tree topology (Upper Guinea-UG (Northern Lower Guinea-LG-North, Southern Lower Guinea-LG-South) (S4 A). In addition to a model with symmetric migration among the three clusters, we fitted a model with no migration allowed between UG and LG-South after the split. For *S. zenkeri*, absent in Upper Guinea, we estimated split times based on the tree topology (LG-North, (LG-South1 and LG-South2)). The observed SFS was compared to the expected SFS under the Isolation with Migration (IM) model with symmetric gene flow (S4 B-E). Using Maximum Likelihood, split time parameters values were estimated in generations. We assumed a mutation rate of μ = 2.5×10 − 9 (1.7×10 − 9 to 3.5×10 − 9) per site per year, estimated for *Populus* (46, 47) and a generation time of 100 years (12, 48). To avoid biasing the demographic inferences due to uneven depth of coverage, which is typical of GBS data, we excluded all singletons, i.e. alleles found only once among the populations, while estimating the demographic parameters. Further, while estimating SFS, admixed individuals with less than 70% genetic ancestry to a single group were excluded.

For each such demographic model, we ran 100 replicates, and selected the parameter estimates from the best fitting model, i.e. the model with the highest log-likelihood, with one important exception. Model fits in δαδl which masked more than 5% of the SFS were excluded. Finally, we performed 1000 replicates for *D. benthamianus*, as this species yielded fewer SNPs and larger sample sizes, in order to get a stable estimate of the demography.

### Gradients of genetic diversity over space and climatic niche modelling

In order to visualise the patterns of genetic diversity over space in Lower Guinea, for each species we calculated the genetic diversity of each of the genotyped individuals, and plotted it against the geographical distance from the closest Last Glacial Maxima refugia (LGM). A mixed effect model was performed using the genetic diversity as the independent variable, the distance to LGM refugia as explanatory variable (minimal distance between the sample and the limit of a postulated refugia) and the genetic cluster as a random variable in order to account for genetic diversity differences across genetic lineages

We calculated the observed heterozygosity (H_o_) of each individual as a proxy for genetic diversity using GenAlEx (49). Admixed individuals with less than 70% genetic ancestry to a single group were excluded. Three different hypotheses of LGM forest refugia were considered. The forest refugia postulated by Maley and Anhuf, based on palaeoclimatic and palynological data (Figure 2): (i) LGM-Maley (14), and (ii) LGM-Anhuf, (27). In addition, specific forest refugia based on the LGM niche models of each species were tested (Figure 2, S7): (iii) LGM-species niche models (see below).

For *P. elata* and *E. suaveolens*, that exhibited inland populations that are equally likely to have survived either in the coastal refugia of Cameroon and Gabon or in the inland refugia in Congo, two additional mixed effects models were run: (iv) LGM-Maley-Coast, including the coastal Maley refugia only, and (v) LGM-Maley-Congo, including the Congo refugia only.

### Species distribution models for the LGM

We collected occurrence records for each species from RAINBIO database (50) and our own records. We visually inspected records to eliminate outliers and deduplicated records per species to obtain one per pixel according to the climate layers resolution (see below). After this, we had 1,029 occurrences distributed among species as follows: *D. benthamianus* (n=427), *E. ivorense* (n=74), *E. suaveolens* (n = 180), *P. elata* (n=118), *S. zenkeri* (n=230) (S7).

We used the 19 bioclim layers from the Worldclim dataset v1.4 at 2.5 arc-min (51) representing the climate between 1960-1990 and calculated the total precipitation of the Austral Summer (December, January and February) using the monthly layers. After performing correlation analysis and Principal component analysis (PCA) we finally selected 5 variables that were minimally correlated to run the models: annual mean temperature (bio1), temperature annual range (bio7), total annual precipitation (bio12), precipitation of the driest month (bio14), and precipitation of December, January and February (pp_djf). We also used the selected variables from simulations with the CCSM4 global climate model for the LGM from Worldclim to project species distributions for that time.

We modelled the distribution of the five species with the biomod2 package in R (52). We randomly selected 5000 pseudoabsences. We used three different algorithms (generalized linear models, GLM; random forest, RF and Maxent) and five repetitions, obtaining a total of 15 models for each species. Models were calibrated using 80% of the occurrences and evaluated using the remaining 20% of occurrences and the TSS and ROC statistics (53). We used a consensus approach to produce an ensemble model using the weighted mean of all models which had at least TSS values above 0.7 and ROC values above 0.8. Binary maps (presence/absence maps) were produced using the TSS threshold (S7).

## ACKNOWLEDGEMENTS

We would like to thank Boris Demenou, Armel Donkpegan, Myriam Heuertz, Théophile Ayolle, Charlotte Hansen, Esra Kaymak, Jean-Louis Doucet, Jerôme Duminil, Jerôme Chave, and Hermann Daniel. This work received financial support from the Marie Curie FP7-PEOPLE-2012-IEF program (project AGORA) awarded to Rosalía Piñeiro.

## REFERENCES

1. Salzmann U & Hoelzmann P (2005) The Dahomey Gap: an abrupt climatically induced rain forest fragmentation in West Africa during the late Holocene. The Holocene 15(2):190–199.

2. Miller CS & Gosling WD (2014) Quaternary forest associations in lowland tropical West Africa. Quaternary Science Reviews 84:7–25.

3. Hey J (2009) The divergence of chimpanzee species and subspecies as revealed in multipopulation isolation-with-migration analyses. Molecular biology and evolution 27(4):921–933.

4. Telfer PT, et al. (2003) Molecular evidence for deep phylogenetic divergence in Mandrillus sphinx. Molecular Ecology 12(7):2019–2024.

5. Anthony NM, et al. (2007) The role of Pleistocene refugia and rivers in shaping gorilla genetic diversity in central Africa. Proceedings of the National Academy of Sciences 104(51):20432–20436.

6. Hardy OJ, et al. (2013) Comparative phylogeography of African rain forest trees: a review of genetic signatures of vegetation history in the Guineo-Congolian region. Comptes Rendus Geoscience 345(7–8):284–296.

7. Heuertz M, Duminil J, Dauby G, Savolainen V, & Hardy OJ (2014) Comparative phylogeography in rainforest trees from Lower Guinea, Africa. PloS one 9(1):e84307.

8. Droissart V, et al. (2018) Beyond trees: biogeographical regionalization of tropical Africa. Journal of biogeography 45(5):1153–1167.

9. Gibbard P, Ehlers J, & Hughes P (2016) Quaternary Glaciations. International Encyclopedia of Geography: People, the Earth, Environment and Technology: People, the Earth, Environment and Technology:1–10.

10. Duminil J, et al. (2015) Late Pleistocene molecular dating of past population fragmentation and demographic changes in African rain forest tree species supports the forest refuge hypothesis. Journal of biogeography 42(8):1443–1454.

11. Faye A, et al. (2016) Phylogenetics and diversification history of African rattans (Calamoideae, Ancistrophyllinae). Botanical journal of the Linnean Society 182(2):256–271.

12. Piñeiro R, Dauby G, Kaymak E, & Hardy OJ (2017) Pleistocene population expansions of shade-tolerant trees indicate fragmentation of the African rainforest during the Ice Ages. Proc. R. Soc. B 284(1866):20171800.

13. Migliore J, et al. (2019) Pre-Pleistocene origin of phylogeographical breaks in African rain forest trees: New insights from Greenwayodendron (Annonaceae) phylogenomics. Journal of biogeography 46(1):212–223.

14. Maley J (1996) The African rain forest–main characteristics of changes in vegetation and climate from the Upper Cretaceous to the Quaternary. Proceedings of the Royal Society of Edinburgh, Section B: Biological Sciences 104:31–73.

15. Hardy OJ, et al. (2019) Seed and pollen dispersal distances in two African legume timber trees and their reproductive potential under selective logging. Molecular ecology 28:3119–3134.

16. White F (1983) The vegetation of Africa: a descriptive memoir to accompany the UNESCO/AETFAT/UNSO vegetation map of Africa by F White. Natural Resources Research Report XX, UNESCO, Paris, France.

17. Fayolle A, et al. (2014) Patterns of tree species composition across tropical African forests. Journal of Biogeography 41(12):2320–2331.

18. Linder HP, et al. (2012) The partitioning of Africa: statistically defined biogeographical regions in sub-Saharan Africa. Journal of Biogeography 39(7):1189–1205.

19. Altschul SF, et al. (1997) Gapped BLAST and PSI-BLAST: a new generation of protein database search programs. Nucleic acids research 25(17):3389–3402.

20. Gagneux P, Gonder MK, Goldberg TL, & Morin PA (2001) Gene flow in wild chimpanzee populations: what genetic data tell us about chimpanzee movement over space and time. Philosophical Transactions of the Royal Society of London. Series B: Biological Sciences 356(1410):889–897.

21. Gonder MK, et al. (1997) A new west African chimpanzee subspecies? Nature 388(6640):337.

22. Fuchs J & Bowie RC (2015) Concordant genetic structure in two species of woodpecker distributed across the primary West African biogeographic barriers. Molecular phylogenetics and evolution 88:64–74.

23. Beresford P & Cracraft J (1999) Speciation in African forest robins (Stiphronis): species limits, phylogentic relationships, and molecular biology. American Museum novitates; no. 3270.

24. Schmidt BK, Foster JT, Angehr GR, Durrant KL, & Fleischer RC (2008) A new species of African forest robin from Gabon (Passeriformes: Muscicapidae: Stiphrornis).

25. Marks BD (2010) Are lowland rainforests really evolutionary museums? Phylogeography of the green hylia (Hylia prasina) in the Afrotropics. Molecular Phylogenetics and Evolution 55(1):178–184.

26. Nicolas V, et al. (2011) The roles of rivers and Pleistocene refugia in shaping genetic diversity in Praomys misonnei in tropical Africa. Journal of Biogeography 38(1):191–207.

27. Anhuf D, et al. (2006) Paleo-environmental change in Amazonian and African rainforest during the LGM. Palaeogeography, Palaeoclimatology, Palaeoecology 239(3–4):510–527.

28. Meunier Q, Moumbogou C, & Doucet J-L (2015) Les arbres utiles du Gabon (Presses agronomiques de Gembloux).

29. Gorel A-P, Fayolle A, & Doucet J-L (2015) Ecology and management of the multipurpose Erythrophleum species (Fabaceae-Caesalpinioideae) in Africa. A review. Biotechnologie, Agronomie, Société et Environnement 19(4):415–429.

30. Duminil J, et al. (2010) CpDNA-based species identification and phylogeography: application to African tropical tree species. Molecular ecology 19(24):5469–5483.

31. Abanda G (2011) La Surexploitation du Scorodophloeus zenkeri Harms (Arbre aAil): Atténuer l Impact de la Gestion Non Durable de l Arbre aAil dans le Massif Forestier de Ngovayang du Sud Cameroun. Editions Universitaires Europeennes.

32. Bruneau A, Mercure M, Lewis GP, & Herendeen PS (2008) Phylogenetic patterns and diversification in the caesalpinioid legumes. Botany 86(7):697–718.

33. Azani N, et al. (2017) A new subfamily classification of the Leguminosae based on a taxonomically comprehensive phylogeny The Legume Phylogeny Working Group (LPWG). Taxon 66(1):44–77.

34. Elshire RJ, et al. (2011) A robust, simple genotyping-by-sequencing (GBS) approach for high diversity species. PloS one 6(5):e19379.

35. Lu F, et al. (2013) Switchgrass genomic diversity, ploidy, and evolution: novel insights from a network-based SNP discovery protocol. PLoS genetics 9(1):e1003215.

36. Bradbury PJ, et al. (2007) TASSEL: software for association mapping of complex traits in diverse samples. Bioinformatics 23(19):2633–2635.

37. Nielsen R, Paul JS, Albrechtsen A, & Song YS (2011) Genotype and SNP calling from next-generation sequencing data. Nature Reviews Genetics 12(6):443.

38. Korneliussen TS, Albrechtsen A, & Nielsen R (2014) ANGSD: analysis of next generation sequencing data. BMC bioinformatics 15(1):356.

39. Schubert M, et al. (2014) Characterization of ancient and modern genomes by SNP detection and phylogenomic and metagenomic analysis using PALEOMIX. Nature protocols 9(5):1056.

40. Lindgreen S (2012) AdapterRemoval: easy cleaning of next-generation sequencing reads. BMC research notes 5(1):337.

41. Li H & Durbin R (2009) Fast and accurate short read alignment with Burrows–Wheeler transform. bioinformatics 25(14):1754–1760.

42. Alexander DH, Novembre J, & Lange K (2009) Fast model-based estimation of ancestry in unrelated individuals. Genome research.

43. Wang J (2017) The computer program STRUCTURE for assigning individuals to populations: easy to use but easier to misuse. Molecular ecology resources 17(5):981–990.

44. Stamatakis A (2014) RAxML version 8: a tool for phylogenetic analysis and post-analysis of large phylogenies. Bioinformatics 30(9):1312–1313.

45. Gutenkunst RN, Hernandez RD, Williamson SH, & Bustamante CD (2009) Inferring the joint demographic history of multiple populations from multidimensional SNP frequency data. PLoS genetics 5(10):e1000695.

46. Tuskan GA, et al. (2006) The genome of black cottonwood, Populus trichocarpa (Torr. & Gray). science 313(5793):1596–1604.

47. Ingvarsson PK (2008) Multilocus patterns of nucleotide polymorphism and the demographic history of Populus tremula. Genetics.

48. Baker TR, et al. (2014) Fast demographic traits promote high diversification rates of Amazonian trees. Ecology Letters 17(5):527–536.

49. Peakall R & Smouse PE (2006) GENALEX 6: genetic analysis in Excel. Population genetic software for teaching and research. Molecular ecology notes 6(1):288–295.

50. Dauby G, et al. (2016) RAINBIO: a mega-database of tropical African vascular plants distributions. PhytoKeys (74):1.

51. Hijmans RJ, Cameron SE, Parra JL, Jones PG, & Jarvis A (2005) Very high resolution interpolated climate surfaces for global land areas. International Journal of Climatology: A Journal of the Royal Meteorological Society 25(15):1965–1978.

52. Thuiller W, Lafourcade B, Engler R, & Araújo MB (2009) BIOMOD–a platform for ensemble forecasting of species distributions. Ecography 32(3):369–373.

53. Zou KH, Liu A, Bandos AI, Ohno-Machado L, & Rockette HE (2011) Statistical evaluation of diagnostic performance: topics in ROC analysis (CRC Press).

54. Mayaux P, Bartholomé E, Fritz S, & Belward A (2004) A new land-cover map of Africa for the year 2000. Journal of Biogeography 31(6):861–877.

