## Supplemental Information for "Contrasting dates of rainforest fragmentation in Africa inferred from trees with different dispersal abilities"

S1. Biological traits important for colonisation in *Pericopsis elata, Distemonanthus benthamianus*, *Erythrophleum ivorense, E. suaveolens*, and *Scorodophloeus zenkeri*. The target species belong to different shade tolerance categories (1): (i) Pioneer (P), species require gaps for establishment, (ii) Non-pioneer light-demanding (NPLD), species can establish in shade but need a gap to reach the canopy, and (iii) Shade-tolerant (ST) species that establish and grow in shade (2). Seed dispersal syndromes include ballistic dispersal, animal dispersal, and wind dispersal. In addition, mean dispersal distances, and the magnitude and frequency of long-distance dispersal (LDD) events are available for *D. benthamianus* and the two *Erythrophleum* species. These were achieved by tracking propagule movement using genetic markers through the estimation of the dispersal distribution (flat tailed distributions are indicative of long distance dispersal events (3).

| Species | Light Tolerance | Dispersal syndrome | Dispersal distance | Long Distance Dispersal | Aggregated distribution | Height | Selfing Rate |
|---|---|---|---|---|---|---|---|
| *Pericopsis elata* | LD, LDP | Wind | 210 | Yes | Yes | 45 | 55 |
| *Distemonanthus benthamianus* | LDP | Wind | 70 | Yes | No | <40 | 7 |
| *Erythrophleum ivorense* | LDP | | | | No | | |
| *Erythrophleum suaveolens* | LD | Ballistic<br>Animals | 200 | No | No | 40 | 0 (30% in seedlings) |
| *Scorodophloeus zenkeri* | ST | Ballistic | | No | Yes | <40 | ? |

Dispersal distance: mean seed dispersal distances in meters calculated by tracking propagule movement using genetic markers (3)
Long Distance Dispersal flat tailed the dispersal distribution calculated by tracking propagule movement using genetic markers through are indicative of long distance dispersal events (3)

S2. Sampling sizes, number of reads and number of SNPs of GBS data of *Pericopsis elata, Distemonanthus benthamianus*, *Erythrophleum ivorense, E. suaveolens*, and *Scorodophloeus zenkeri*.

| | Nr. RAD libraries | Nr. samples ingroup | Nr. samples outgroup | Nr. raw reads/ sample | Nr. mapped reads/ sample | Depth coverage | Average length of mapping reads | nr. SNPS TASSEL | nr. SNPS PALEOMIX |
|---|---|---|---|---|---|---|---|---|---|
| *P. elata* | 92 | **45** | 1 | 3387236 | 568240 | 20 | 50 | 10948 | 14904 |
| *D. benthamianus* | 88 | **42** | 2 | 5518179 | 1561119 | 32 | 58 | 10665 | 17921 |
| *Erythrophleum* | 90 | 42 | 3 | 4112432 | 1279269 | 12 | 58 | | |
| *E. ivorense* | | 18 | | | | | | 25534 | 29589 |
| *E. suaveolens* | | 24 | | | | | | 27838 | 61667 |
| *S. zenkeri* | 94 | **46** | 1 | 4969330 | 1500869 | 20 | 56 | 17914 | 70271 |

S3. RAxML phylogenetic analyses of 189 individuals using the GBS genotype calls with outgroups for *Pericopsis elata, Distemonanthus benthamianus*, *Erythrophleum ivorense, E. suaveolens*, and *Scorodophloeus zenkeri.* Branch width is proportional to bootstrap supports.

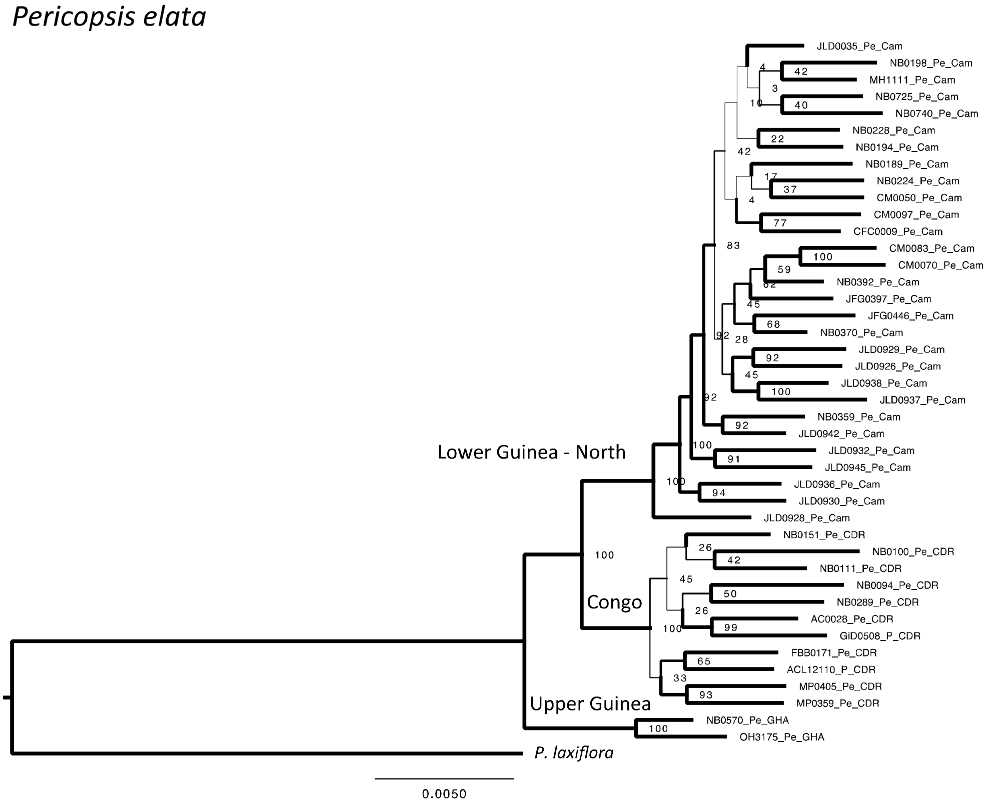


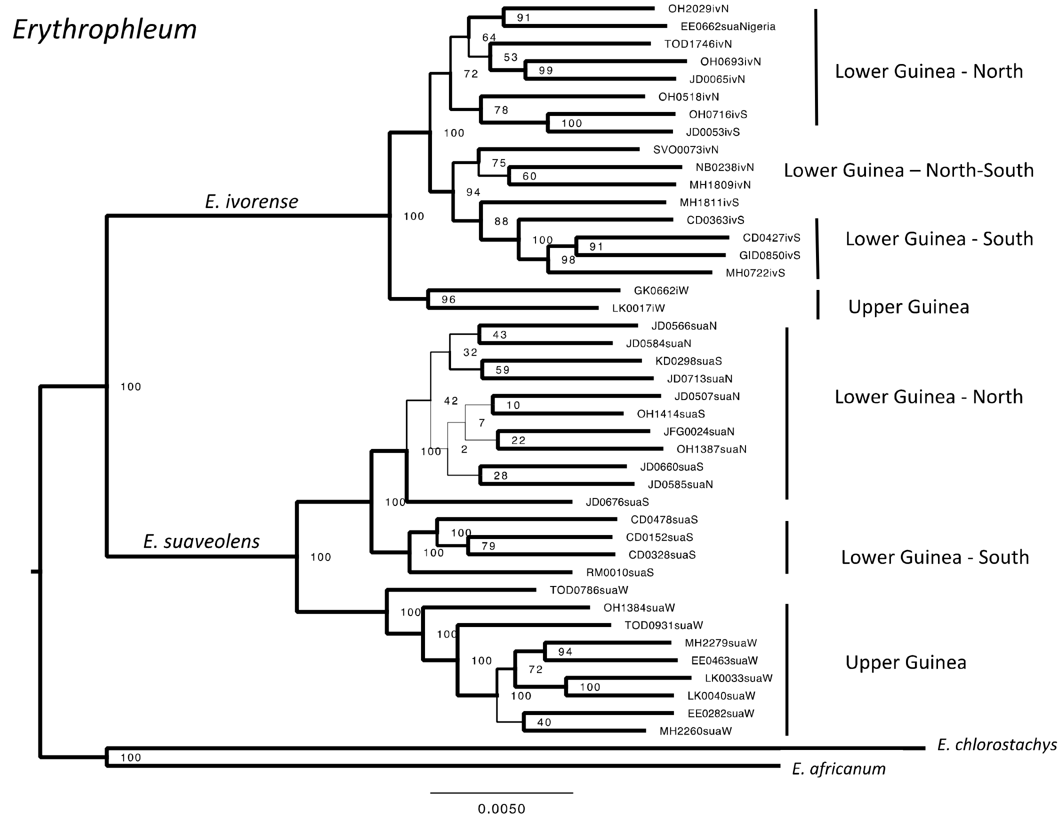

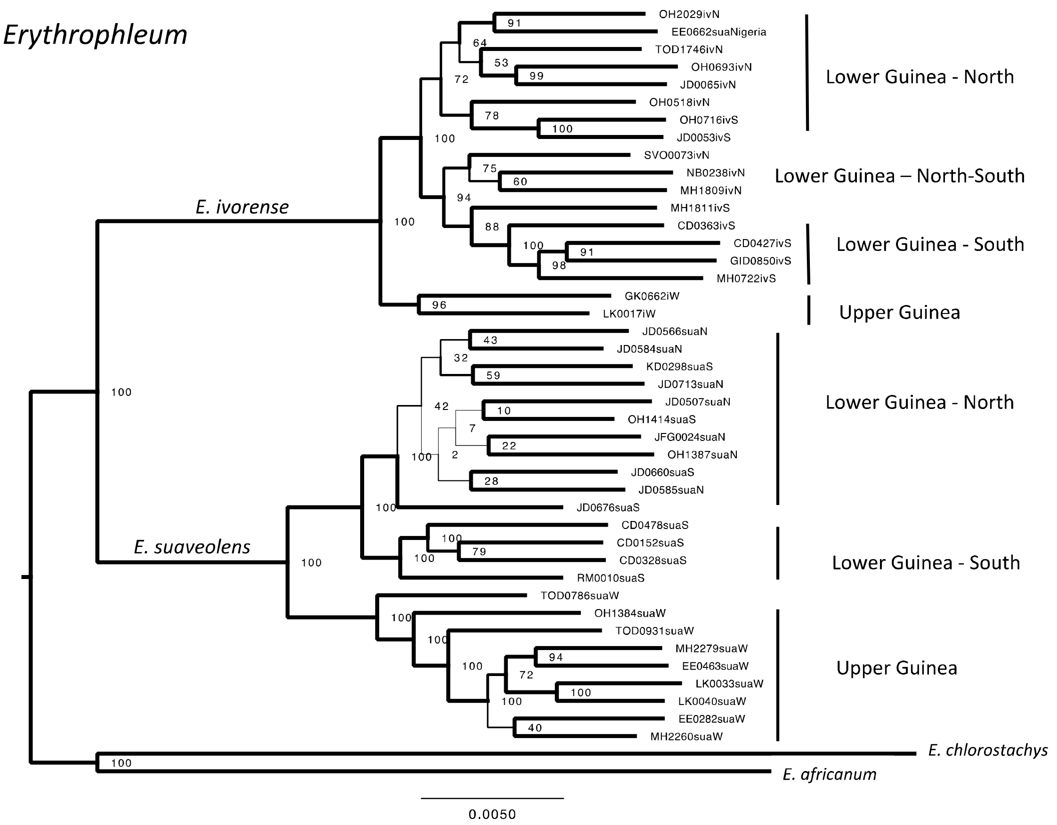
Erythrophleum
E. ivorense
E. suaveolens
OH2029ivN
EE0662suaNigeria
TOD1746ivN
OH0693ivN
JD0065ivN
OH0518ivN
OH0716ivS
JD0053ivS
SVO0073ivN
NB0238ivN
MH1809ivN
MH1811ivS
CD0363ivS
CD0427ivS
GID0850ivS
MH0722ivS
GK0662iW
LK0017iW
JD0566suaN
JD0584suaN
KD0298suaS
JD0713suaN
JD0507suaN
OH1414suaS
JFG0024suaN
OH1387suaN
JD0660suaS
JD0585suaN
JD0676suaS
CD0478suaS
CD0152suaS
CD0328suaS
RM0010suaS
TOD0786suaW
OH1384suaW
TOD0931suaW
MH2279suaW
EE0463suaW
LK0033suaW
LK0040suaW
EE0282suaW
MH2260suaW
E. chlorostachys
E. africanum
Lower Guinea - North
Lower Guinea – North-South
Lower Guinea - South
Upper Guinea
Lower Guinea - North
Lower Guinea - South
Upper Guinea
0.0050

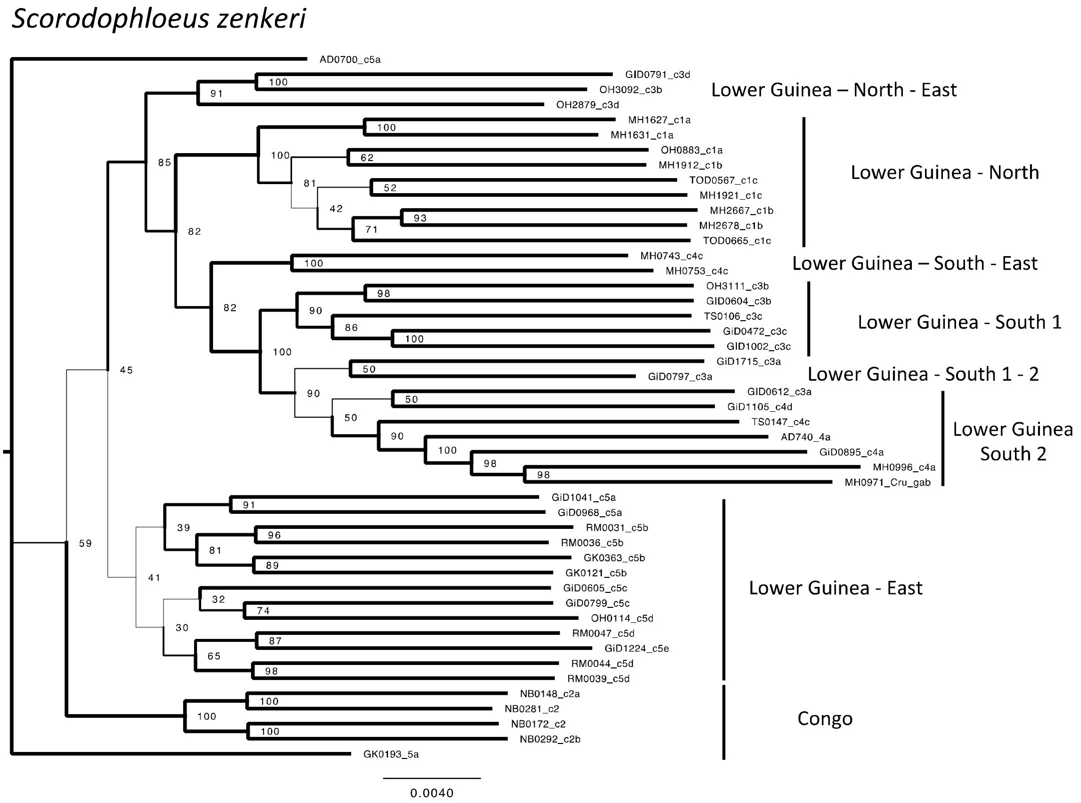
Scorodophloeus zenkeri
AD0700_c5a
GID0791_c3d
OH3092_c3b
OH2879_c3d
MH1627_c1a
MH1631_c1a
OH0883_c1a
MH1912_c1b
TOD0567_c1c
MH1921_c1c
MH2667_c1b
MH2678_c1b
TOD0665_c1c
MH0743_c4c
MH0753_c4c
OH3111_c3b
GID0604_c3b
TS0106_c3c
GiD0472_c3c
GID1002_c3c
GiD1715_c3a
GiD0797_c3a
GID0612_c3a
GiD1105_c4d
TS0147_c4c
AD740_4a
GiD0895_c4a
MH0996_c4a
MH0971_Cru_gab
GiD1041_c5a
GiD0968_c5a
RM0031_c5b
RM0036_c5b
GK0363_c5b
GK0121_c5b
GiD0605_c5c
GiD0799_c5c
OH0114_c5d
RM0047_c5d
GiD1224_c5e
RM0044_c5d
RM0039_c5d
NB0148_c2a
NB0281_c2
NB0172_c2
NB0292_c2b
GK0193_5a
Lower Guinea – North - East
Lower Guinea - North
Lower Guinea – South - East
Lower Guinea - South 1
Lower Guinea - South 1 - 2
Lower Guinea South 2
Lower Guinea - East
Congo
0.0040

S4. Demographic models tested with δαδι (A): Based on the admixture and phylogeny results, for the three species widespread in Upper and Lower Guinea -*Distemonanthus bentamianus*, *Erythrophleum suaveolens*, *E. ivorense*-, we fitted a three-population model with the tree topology (Upper Guinea-UG (Northern Lower Guinea-LG-North, Southern Lower Guinea-LG-South). For *Scorodophleous zenkeri*, absent in Upper Guinea, we estimated split times based on the tree topology (LG-North, (LG-South1 and LG-South2)). For the best-fitting model under Isolation with Migration the following demographic parameters were estimated: Divergence times ($T_1$, $T_2$), Population size ($N_1$, $N_2$, $N_3$, $N_4$, $N_5$), and Migration rate ($m_{12}$, $m_{34}$, $m_{35}$, $m_{45}$). The various parameters fitted are marked on the model. The parameter $m_{35}$ (marked in red), the migration rate between population UG and LG-South, is set to 0, in models where the hypothesis of no migration between Upper Guinea and Lower Guinea-South. (B-E) Fit of the most likely models to the data for *D. benthamianus* (B), *E. ivorense* (C), *E. suaveolens* (D), *S. zenkeri* (E) are also shown. The panels from top to bottom show the observed SFS, the fitted SFS, the difference between the two, and finally the histogram of mismatches between the two SFS

A

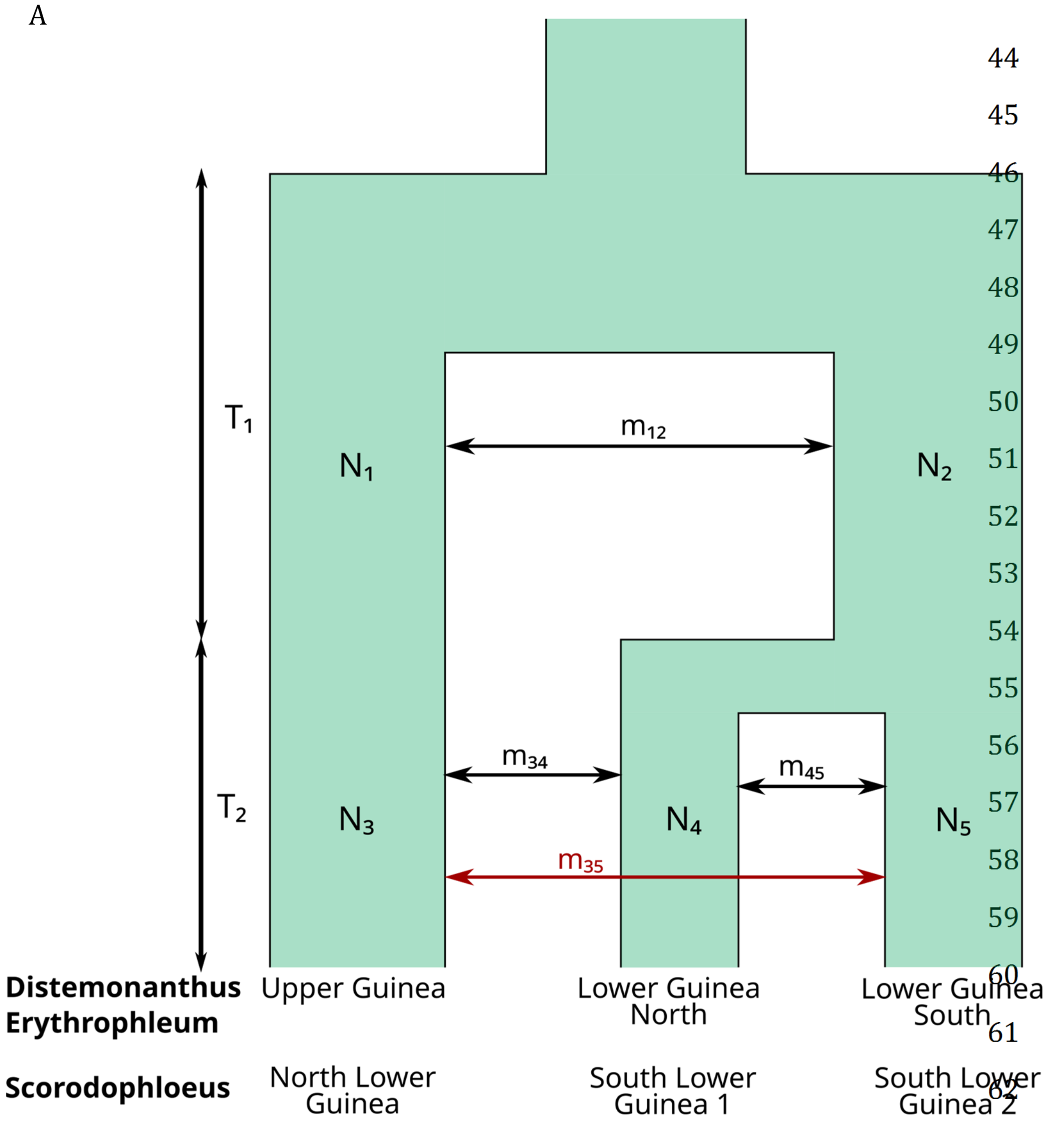

B *Distemonanthus benthamianus*

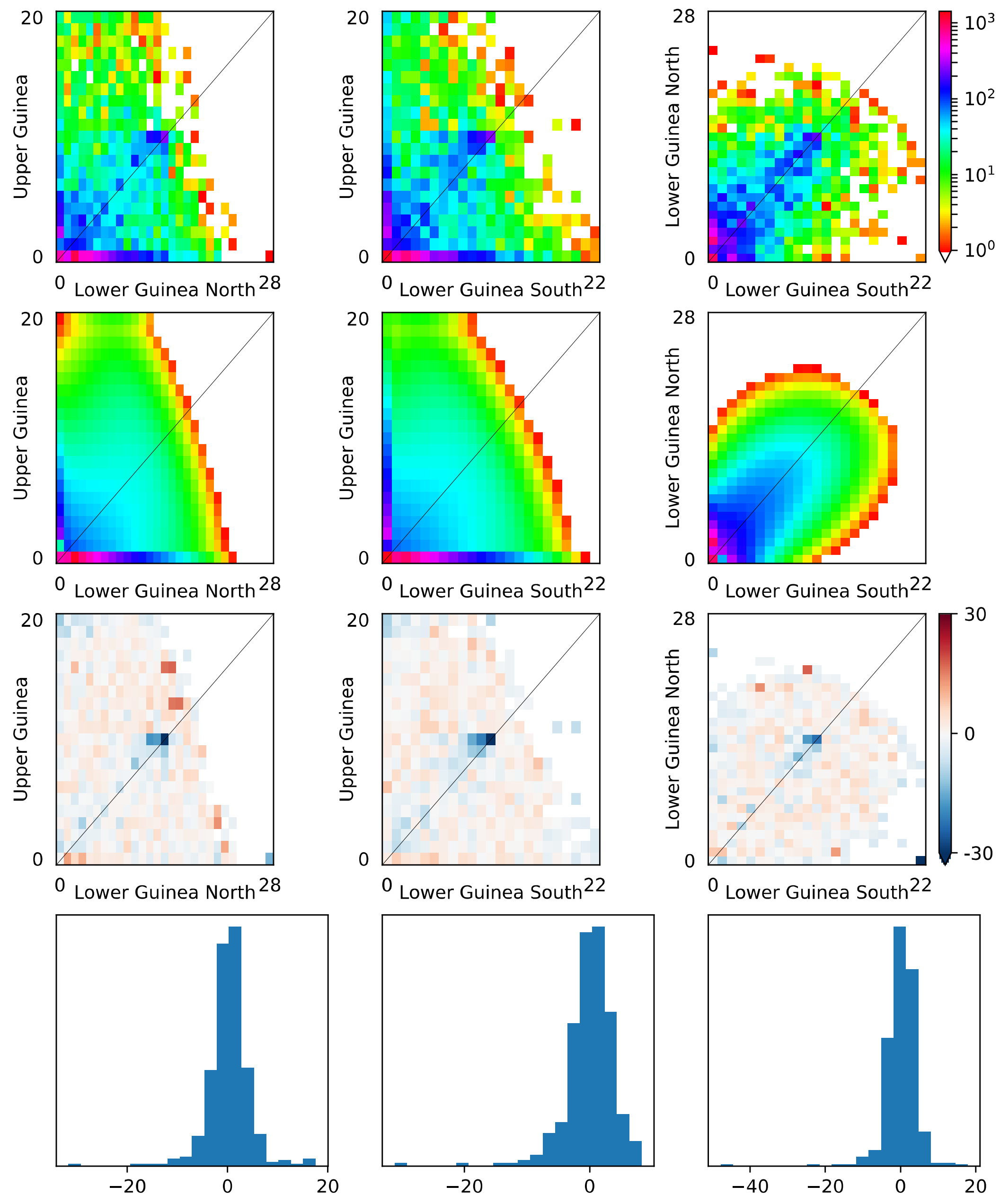

B *Distemonanthus benthamianus*

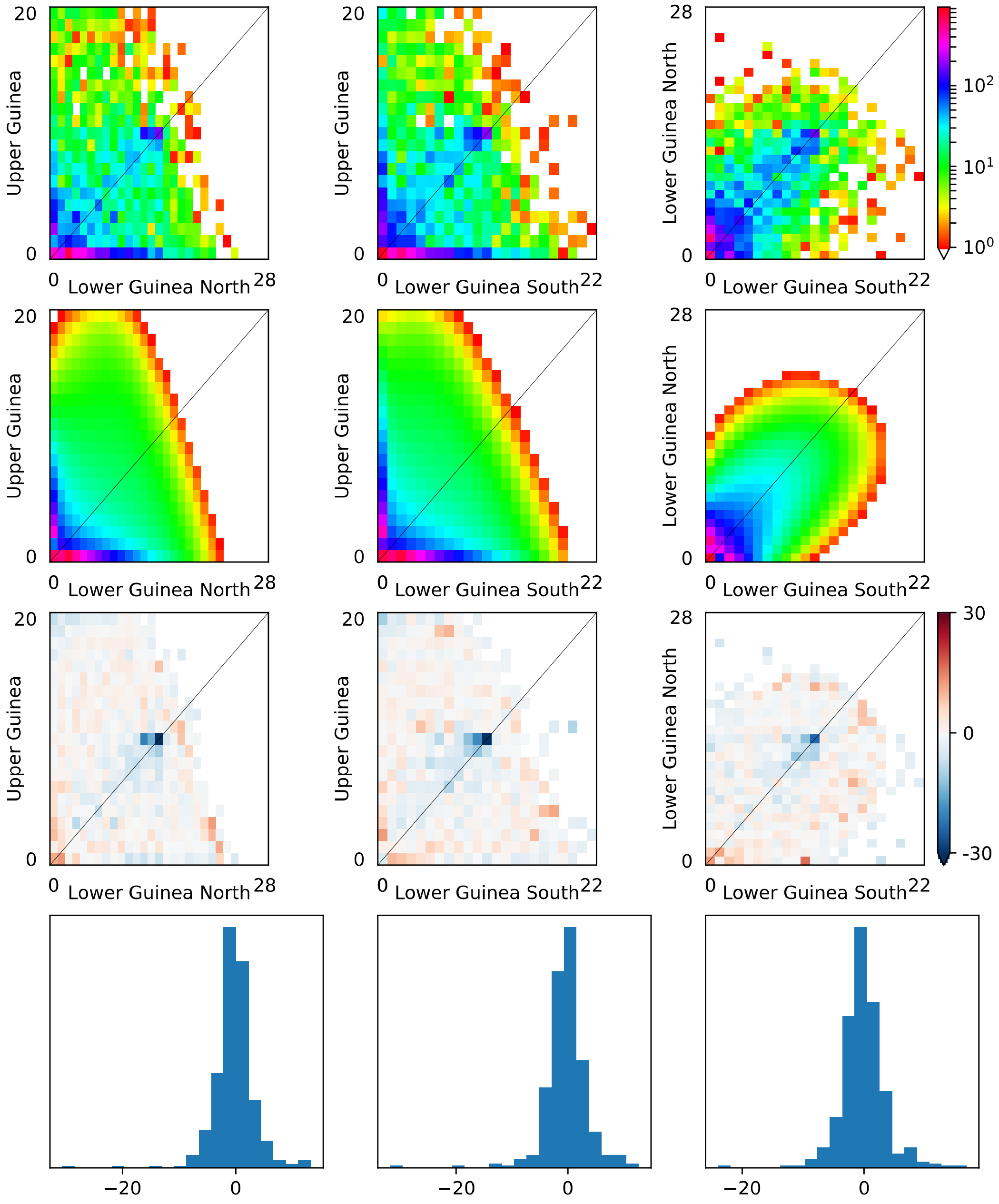

B *Distemonanthus benthamianus*

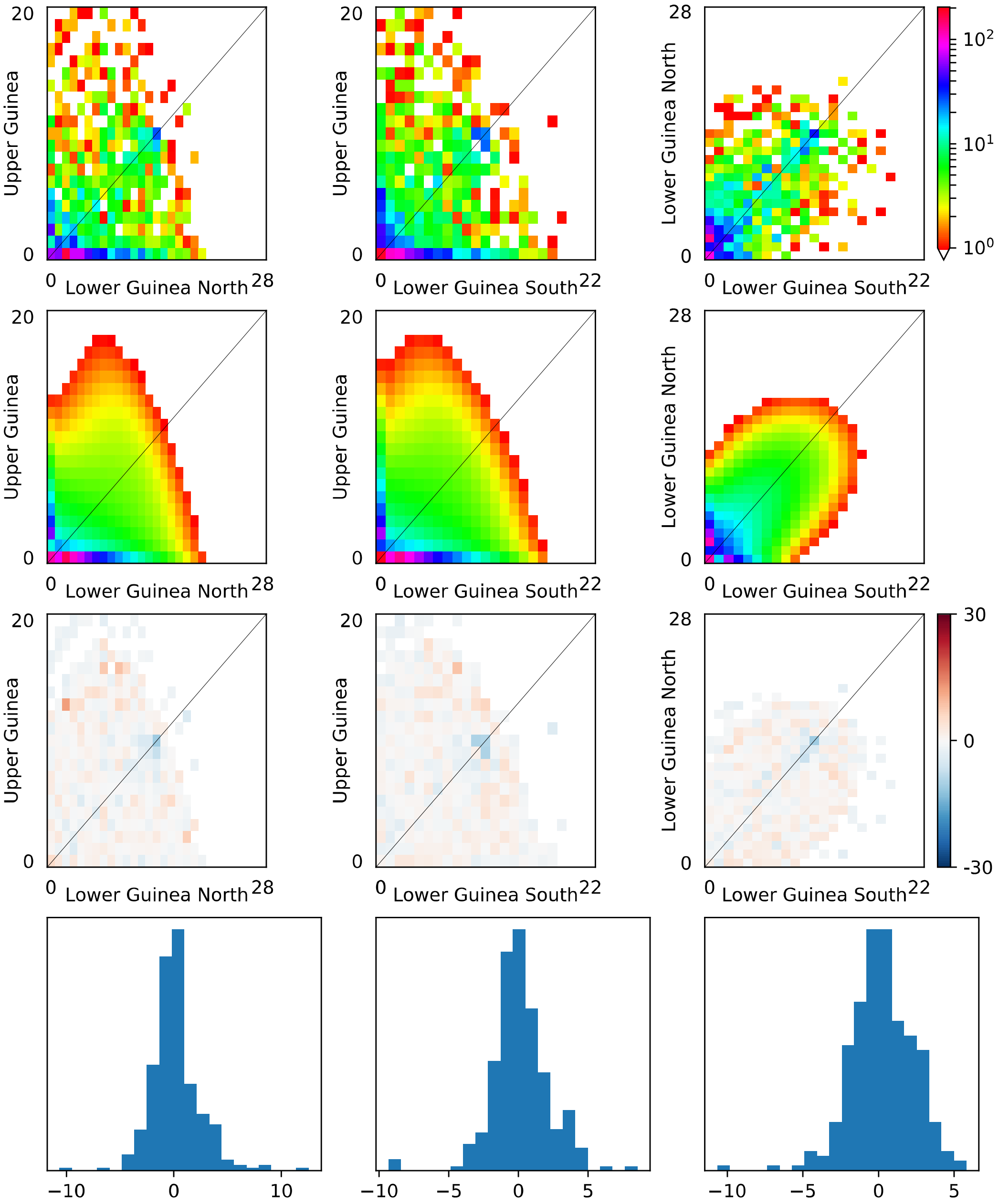

C *Erythrophleum ivorense*

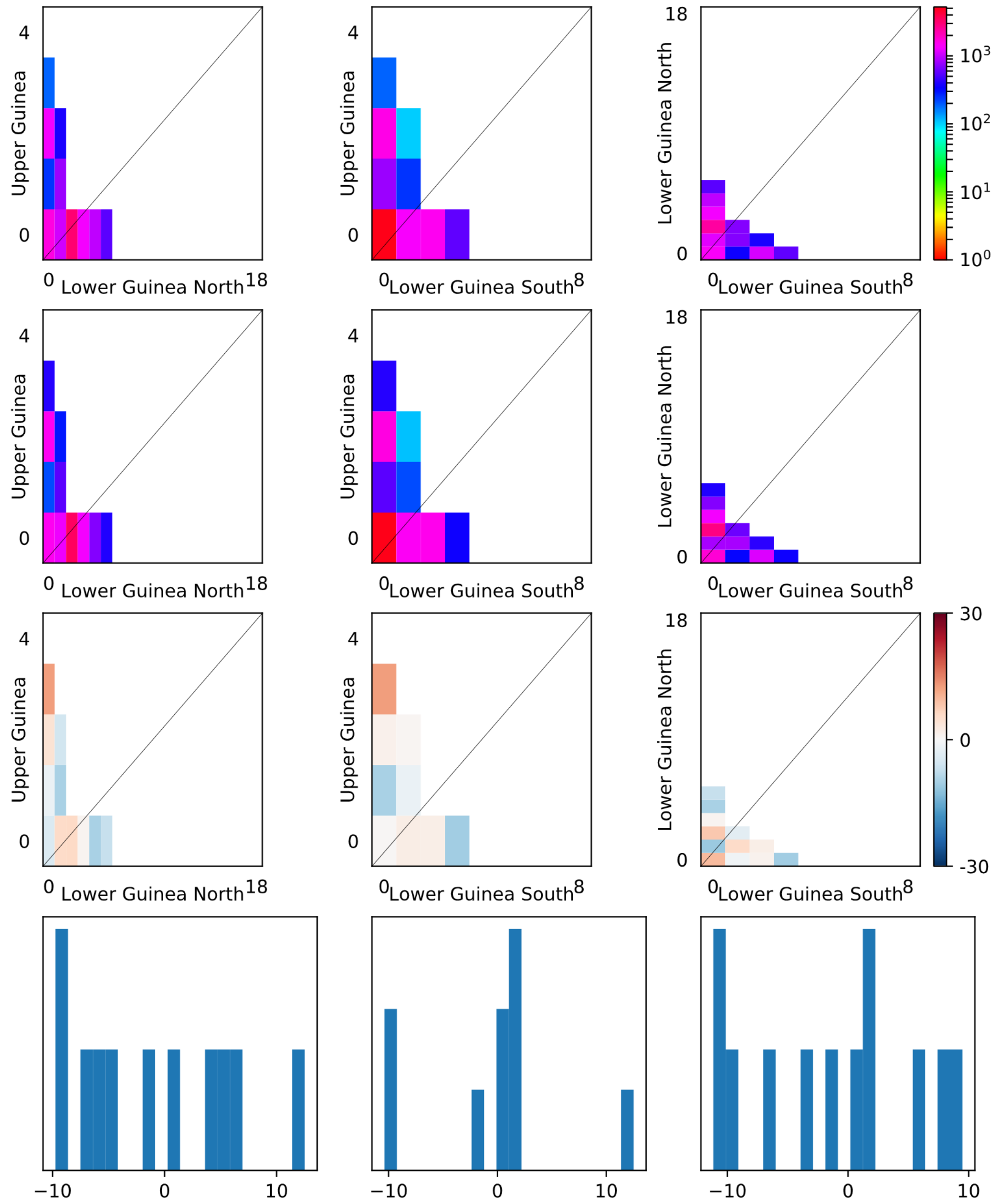

C *Erythrophleum ivorense*

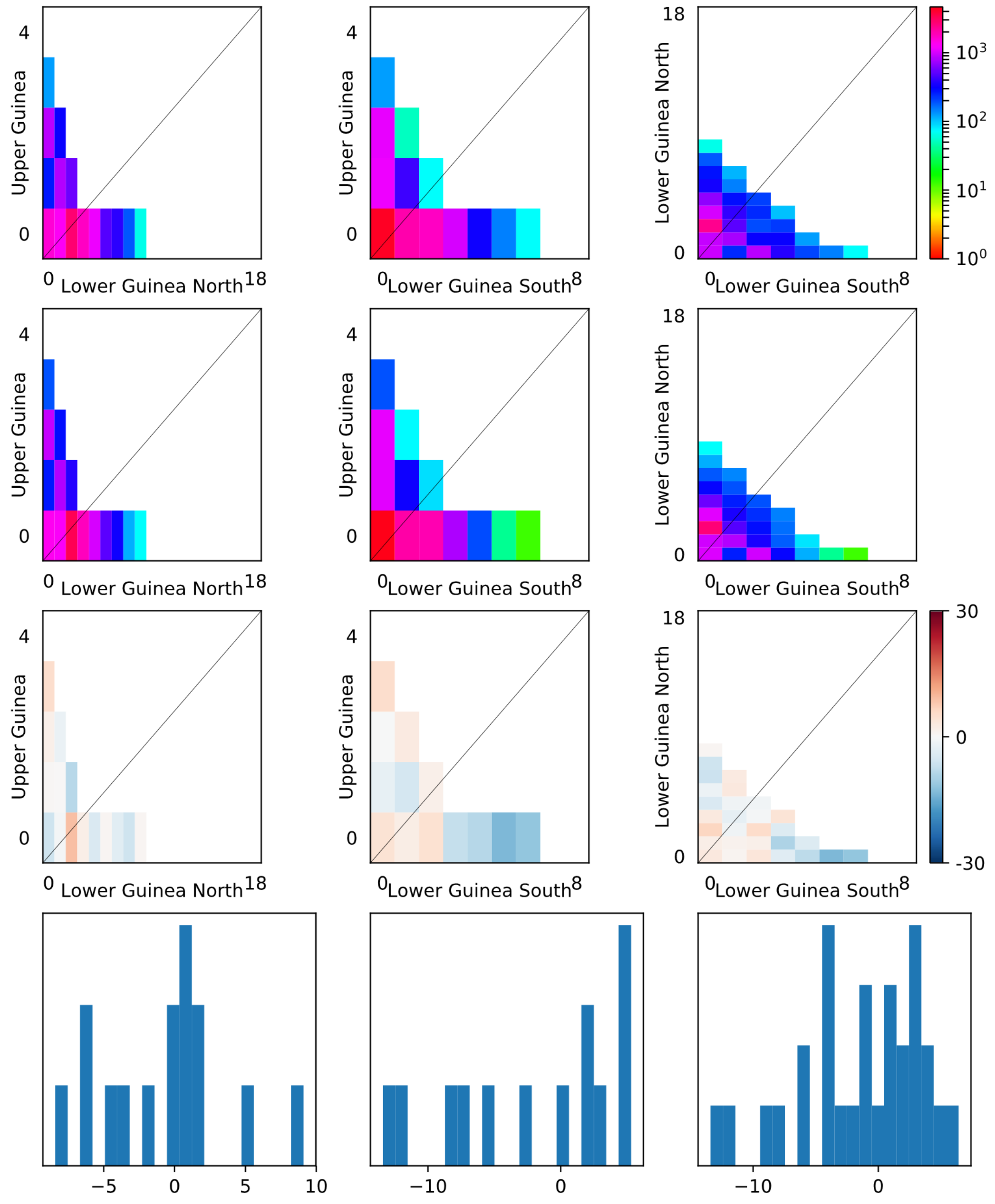


72

73

C *Erythrophleum ivorense*

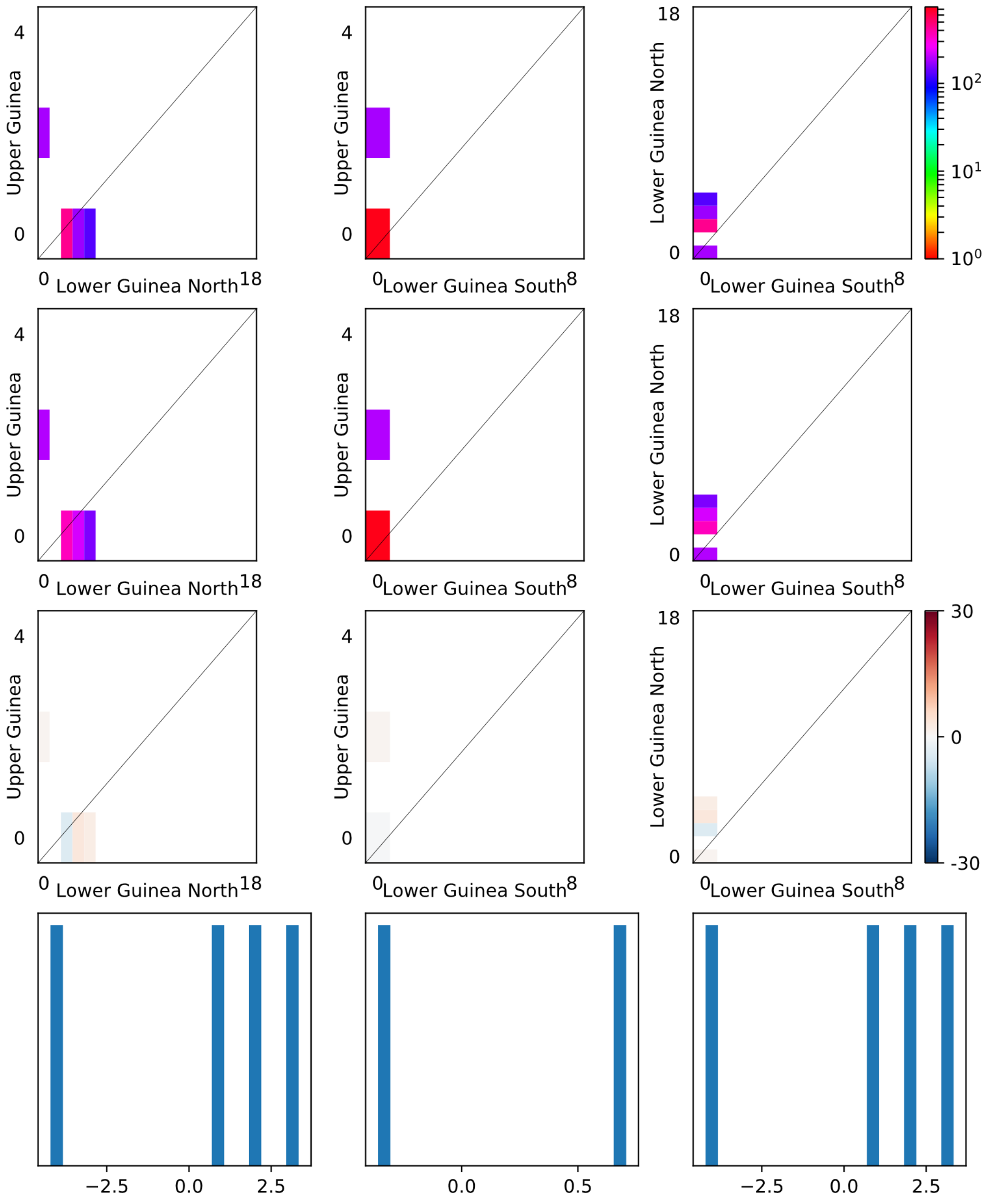

D *Erythrophleum suaveolens*

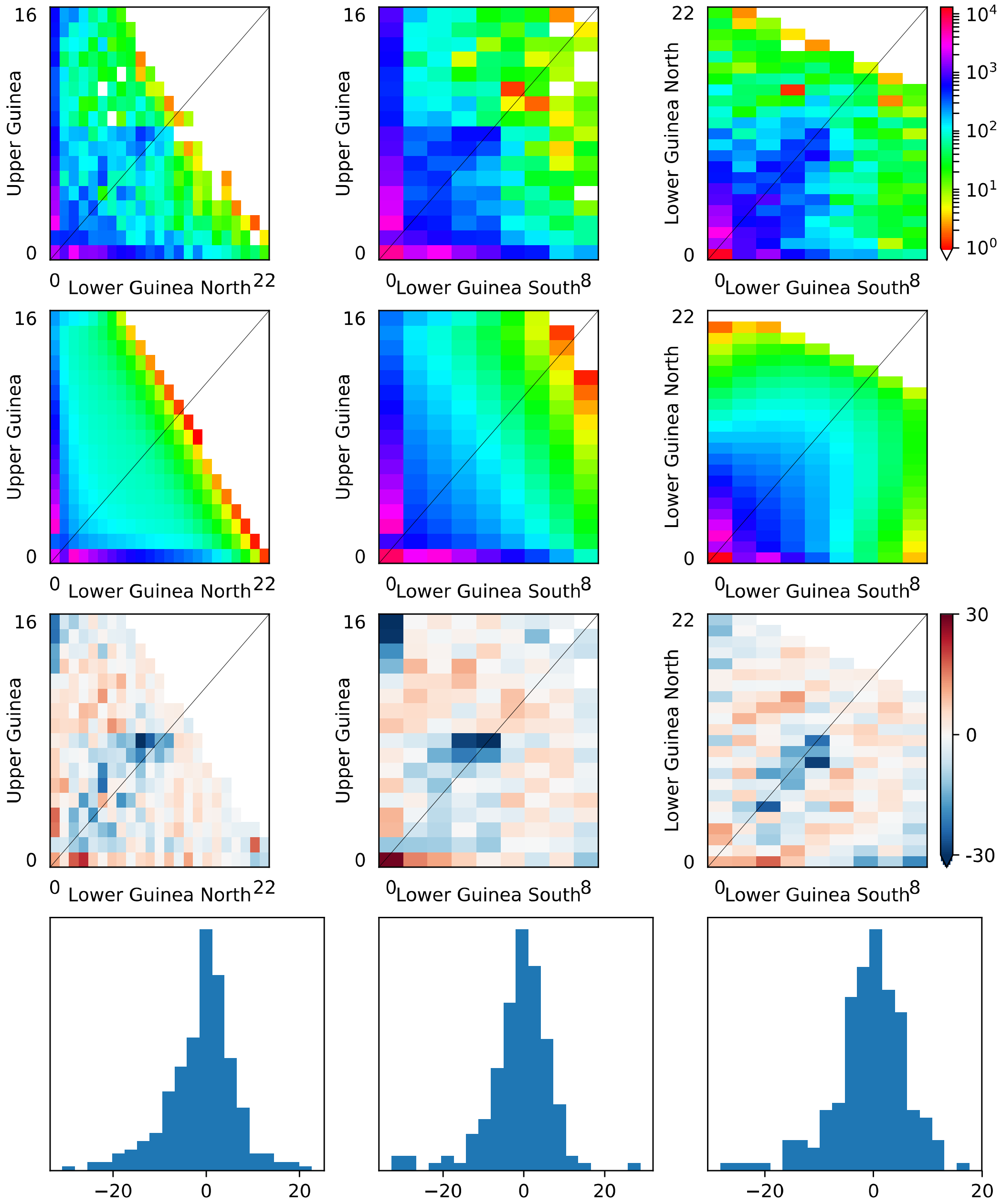


76

77

D *Erythrophleum suaveolens*

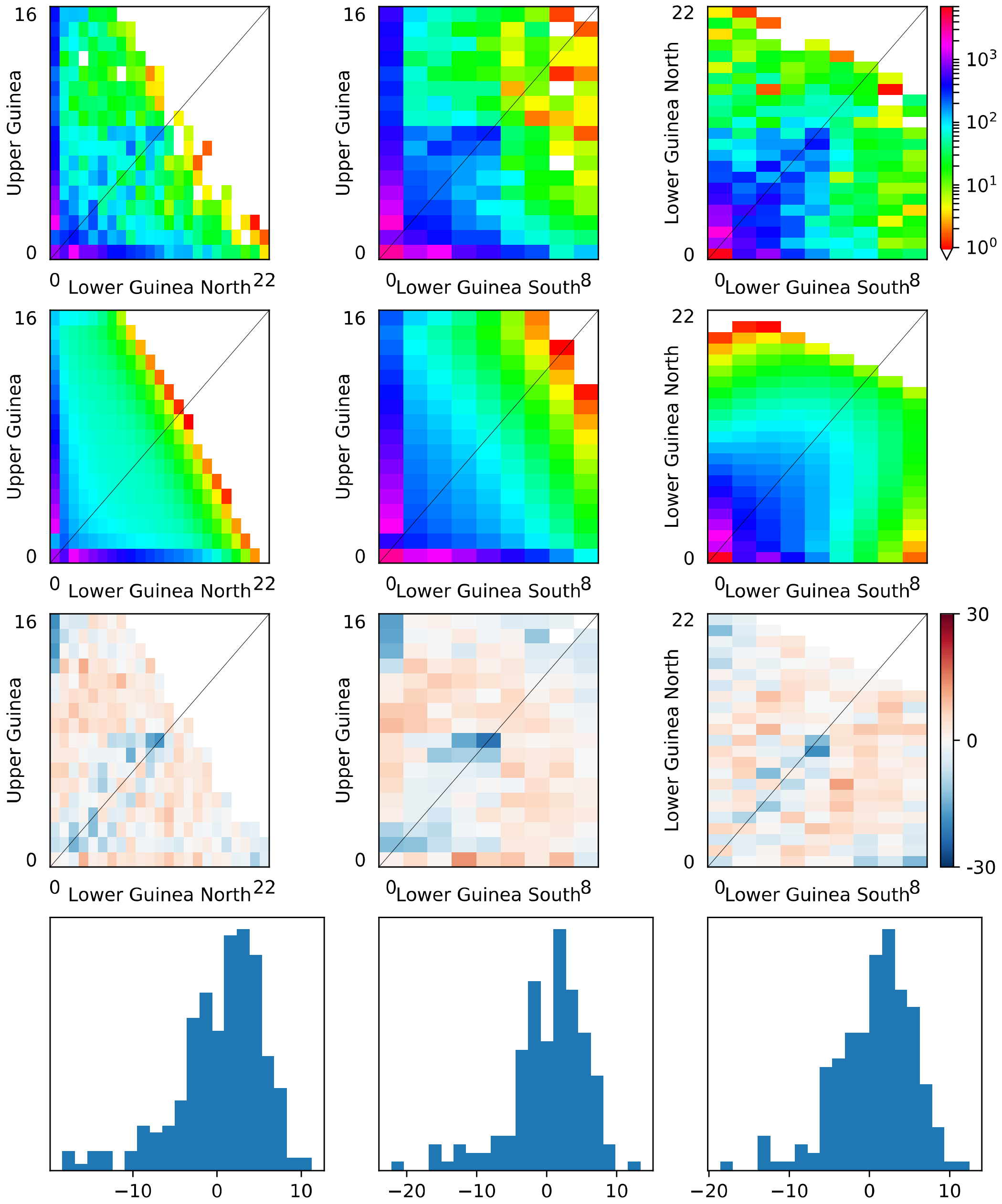


78

79

D *Erythrophleum suaveolens*

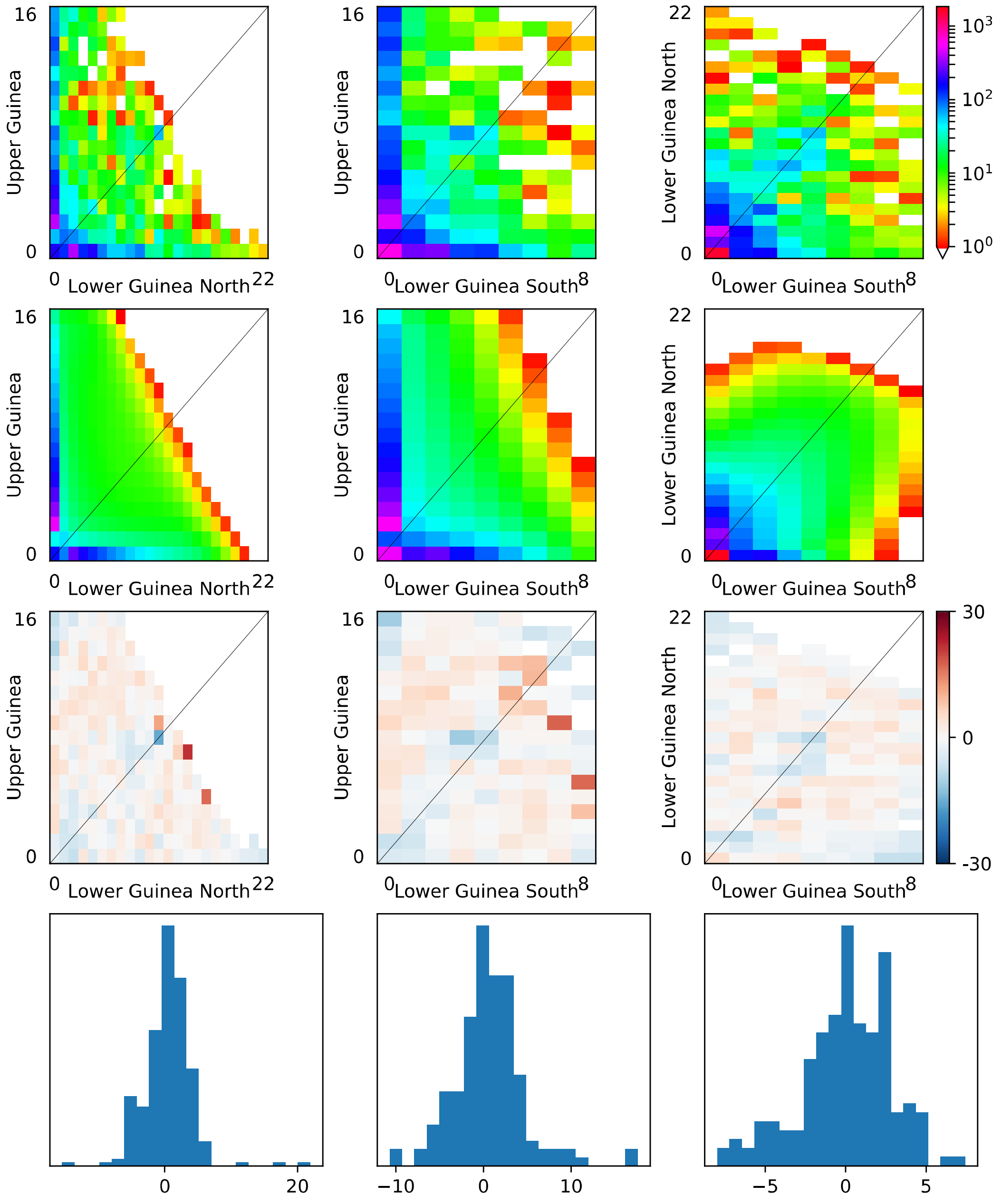


80

81

E *Scorodophloeus zenkeri*

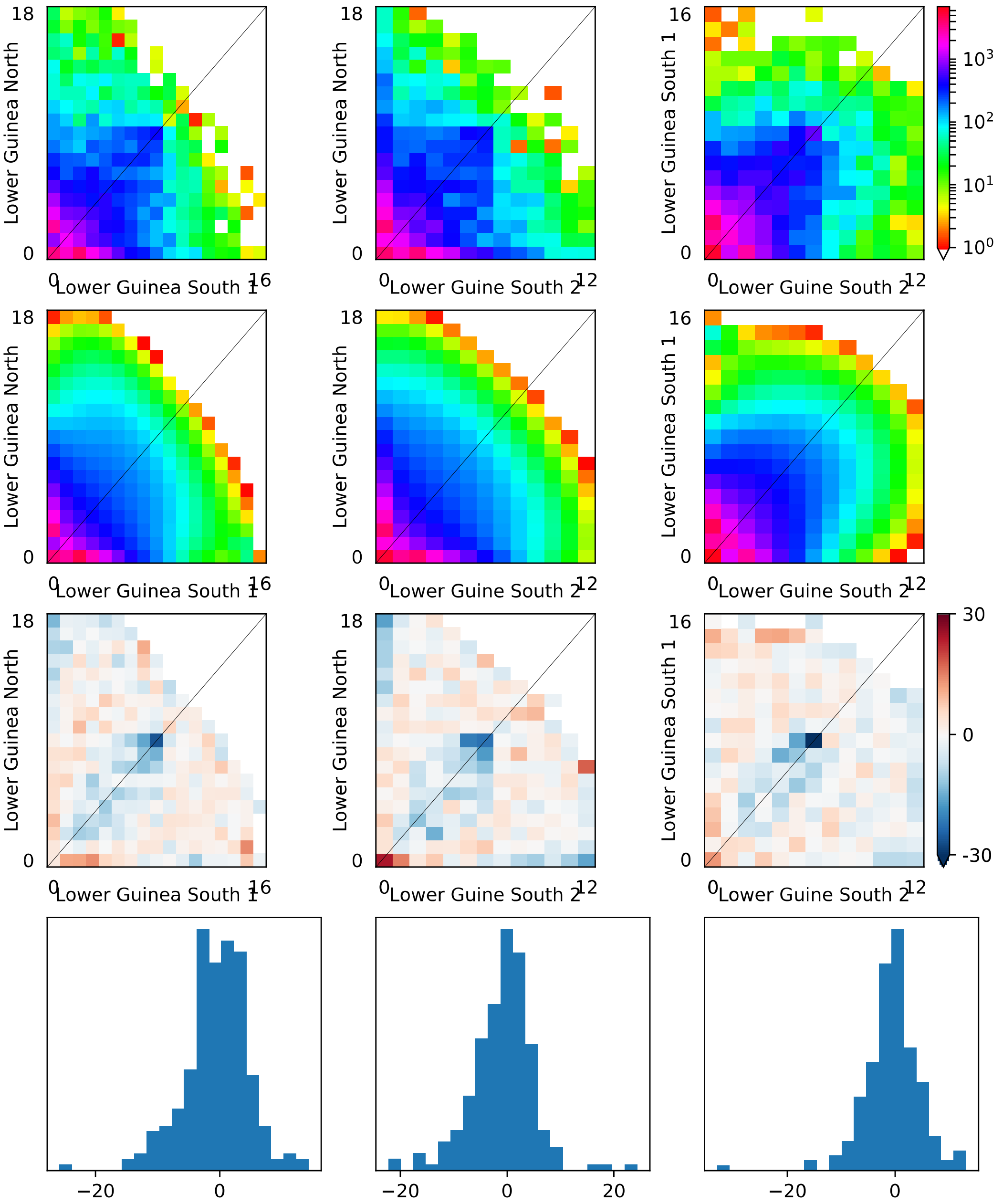


82

83

E *Scorodophloeus zenkeri*

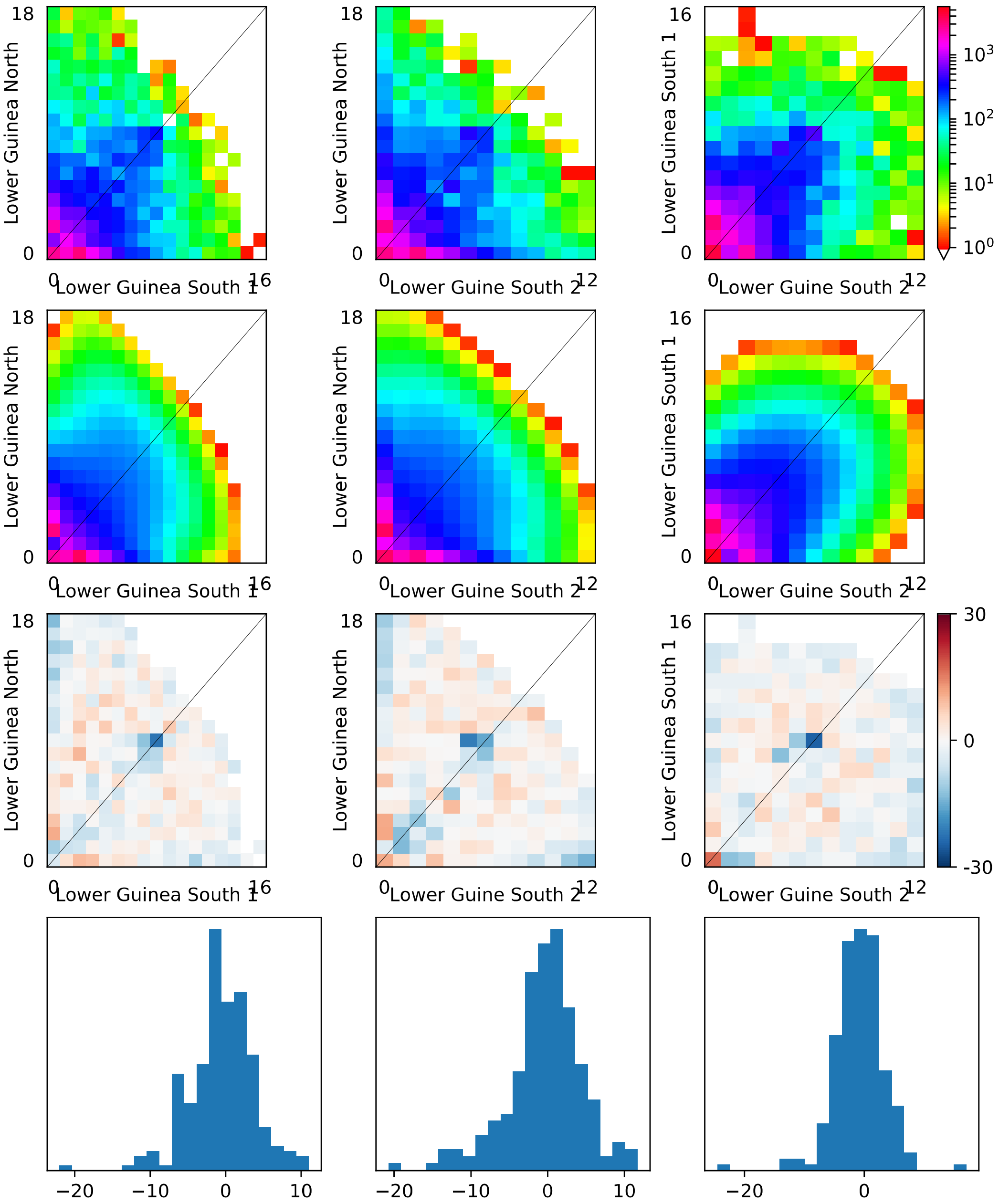

E *Scorodophloeus zenkeri*

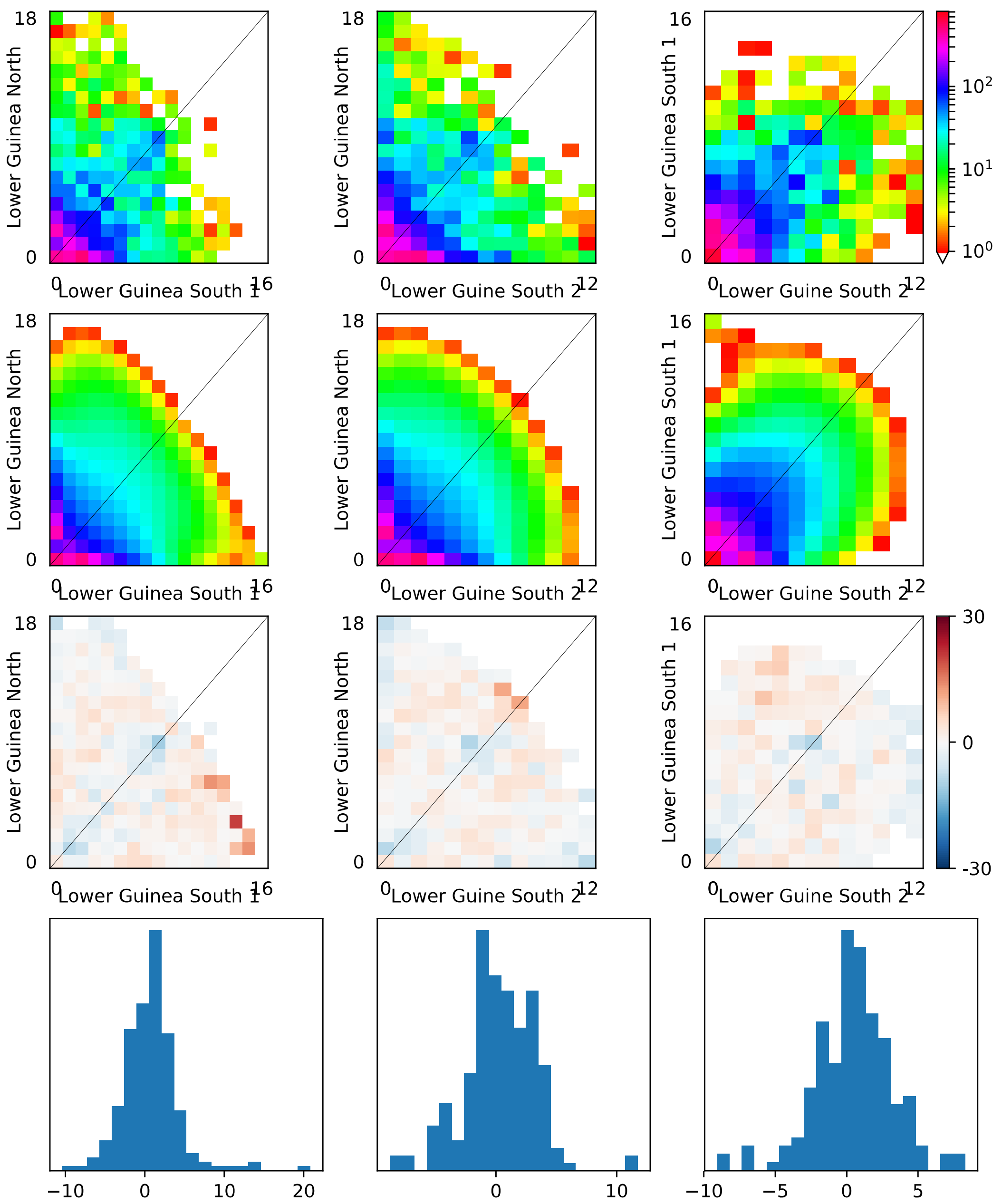

S5. Demographic history of each species using $\partial a \partial$ I: *Distemonanthus benthamianus, Erthrophleum ivorense*, *E. suaveolens, and Scorodophloeus zenkeri* Harms. Demographic parameters (divergence time in years, population size, and gene flow) for the best-fitting model under Isolation with Migration. The estimates are provided under two scenarios, one with continuous migration between all extant populations in Upper Guinea (UG), Lower Guinea-North (nLG), and Lower Guinea South (sLG), and second, a scenario where the migration between UG and sLG ceases after the split into nLG and sLG. Note that this model is only applicable to the three species occurring in Upper Guinea, *E. suaveolens*, *E. ivorense,* and *D. benthamianus*. The demographic parameter estimates are provided for three datasets per species, depending on the stringency of the filtering criteria, filtering 1, the most lenient and filtering 3, the most stringent.

| | CONTINUOUS MIGRATION UG-sLG | | | NO MIGRATION UG-sLG after SPLIT | | | |
|---|---|---|---|---|---|---|---|
| Dataset FILTERING 1 | *D. benthamianus* | *E. ivorense* | *E. suaveolens* | *D. benthamianus* | *E. ivorense* | *E. suaveolens* | *S. zenkeri* |
| Divergence time (years ago) | | | | | | | |
| nLG – sLG | 32762.10 | 101702.00 | 186174.00 | 28982.50 | 103062.00 | 359277.00 | 3436990.00 |
| UG – (nLG, sLG) | 1045800.00 | 1294670.00 | 3848430.00 | 1224790.00 | 1151310.00 | 3493440.00 | |
| sLG1, sLG2 | | | | | | | 1462520.00 |
| Effective Population sizes | | | | | | | |
| nLG | 7264.07 | 2680.88 | 4664.77 | 133153.00 | 6245.23 | 5506.23 | 5258.66 |
| sLG | 1432.23 | 5769.04 | 16354.40 | 1324.96 | 4744.03 | 28493.90 | 8058.34 |
| UG | 806.20 | 5596.17 | 3937.39 | 543.00 | 5530.09 | 5332.62 | |
| sLG2 | | | | | | | 5845.59 |
| Gene flow | | | | | | | |
| nLG – sLG | 5.527E-07 | 4.845E-05 | 1.579E-05 | 5.745E-06 | 3.015E-05 | 1.836E-05 | 2.16E-05 |
| UG – nLG | 3.076E-05 | 2.638E-04 | 5.347E-06 | 1.203E-05 | 1.697E-05 | 1.162E-05 | |
| UG – sLG | 4.630E-06 | 1.476E-05 | 4.181E-06 | | | | |
| UG – (nLG, sLG) | 1.006E-04 | 4.619E-05 | 3.543E-05 | 2.032E-04 | 6.342E-05 | 3.555E-05 | |
| sLG1 – sLG2 | | | | | | | 8.07E-05 |
| nLG – sLG1 | | | | | | | 4.32E-05 |
| nLG – sLG2 | | | | | | | 2.28E-05 |
| Model details | | | | | | | |
| Log-likelihood | -16246.28 | -1222.39 | -25559.17 | -15764.60 | -394.29 | -25466.29 | -17122.81 |
| # of SNPs | 17921.38 | 29589.06 | 61667.59 | 17921.38 | 29513.13 | 61667.59 | 70271.16 |
| Dataset FILTERING 2 | *D. benthamianus* | *E. ivorense* | *E. suaveolens* | *D. benthamianus* | *E. ivorense* | *E. suaveolens* | *S. zenkeri* |

| | | | | | | | |
|---|---|---|---|---|---|---|---|
| Divergence time (years ago) | | | | | | | |
| nLG – sLG | 23035.10 | 140946.00 | 172406.00 | 7698.92 | 76898.30 | 101781.00 | 3085940.00 |
| UG – (nLG, sLG) | 531865.00 | 920277.00 | 3078110.00 | 115531.00 | 1034190.00 | 1053050.00 | |
| sLG1, sLG2 | | | | | | | 174028.00 |
| Effective Population sizes | | | | | | | |
| nLG | 2283.91 | 3856.03 | 6161.07 | 5748.17 | 5831.60 | 2535.06 | 19839.60 |
| sLG | 1846.64 | 12244.50 | 19425.10 | 390.51 | 2928.73 | 3872.18 | 61195.50 |
| UG | 1831.27 | 3073.96 | 5139.19 | 4864.58 | 7311.60 | 1631.48 | |
| sLG2 | | | | | | | 19839.60 |
| Gene flow | | | | | | | |
| nLG – sLG | 1.869E-05 | 2.954E-05 | 3.674E-06 | 6.035E-05 | 2.631E-05 | 4.643E-05 | 8.33E-05 |
| UG – nLG | 1.499E-04 | 1.187E-04 | 9.769E-06 | 4.510E-06 | 4.409E-05 | 3.167E-05 | |
| UG – sLG | 1.606E-07 | 4.544E-05 | 7.888E-06 | | | | |
| UG – (nLG, sLG) | 1.090E-04 | 5.617E-05 | 2.564E-05 | 9.472E-05 | 8.918E-05 | 4.937E-05 | |
| sLG1 – sLG2 | | | | | | | 1.39E-05 |
| nLG – sLG1 | | | | | | | 1.00E-06 |
| nLG – sLG2 | | | | | | | 5.62E-06 |
| Model details | | | | | | | |
| Log-likelihood | -11535.02 | -852.11 | -15264.09 | -11098.82 | -637.26 | -14168.11 | -13590.11 |
| # of SNPs | 11323.50 | 22792.46 | 35042.61 | 11323.50 | 22151.64 | 35042.61 | 54156.53 |
| Dataset FILTERING 3 | *D. benthamianus* | *E. ivorense* | *E. suaveolens* | *D. benthamianus* | *E. ivorense* | *E. suaveolens* | *S. zenkeri* |
| Divergence time (years ago) | | | | | | | |
| nLG – sLG | 303853.00 | 18680.10 | 102121.00 | 14433.40 | 136388.00 | 28257.80 | 576863.00 |
| UG – (nLG, sLG) | 836418.00 | 402973.00 | 989674.00 | 188002.00 | 223053.00 | 1719930.00 | |
| sLG1, sLG2 | | | | | | | 224655.00 |
| Effective Population sizes | | | | | | | |
| nLG | 624.65 | 10567.30 | 1918.23 | 3532.07 | 843.08 | 925.61 | 997.41 |
| sLG | 705.28 | 344.60 | 6127.32 | 829.15 | 523.97 | 1568.86 | 863.55 |
| UG | 421.76 | 6916.96 | 1826.36 | 3816.03 | 667.55 | 12721.40 | |
| sLG2 | | | | | | | 1704.64 |
| Gene flow | | | | | | | |

| | | | | | | | |
|---|---|---|---|---|---|---|---|
| nLG – sLG | 1.323E-03 | 1.001E-05 | 2.091E-05 | 1.029E-04 | 2.752E-05 | 7.251E-07 | 2.51E-04 |
| UG – nLG | 5.980E-04 | 2.728E-04 | 1.966E-05 | 5.153E-06 | 8.005E-05 | 5.126E-05 | |
| UG – sLG | 1.106E-05 | 1.154E-04 | 1.860E-05 | | | | |
| UG – (nLG, sLG) | 1.800E-04 | 1.111E-04 | 7.537E-05 | 5.422E-04 | 4.433E-04 | 7.772E-05 | |
| sLG1 – sLG2 | | | | | | | 5.52E-04 |
| nLG – sLG1 | | | | | | | 2.74E-04 |
| nLG – sLG2 | | | | | | | 8.19E-05 |
| Model details | | | | | | | |
| Log-likelihood | -3445.92 | -53.13 | -4662.02 | -3387.54 | -31.23 | -4420.33 | -4600.70 |
| # of SNPs | 2428.57 | 6093.65 | 7588.75 | 2428.57 | 5395.92 | 7588.75 | 9782.24 |

S6. Decline of genetic diversity, based on Genotyping by Sequencing, with distance from refugia coastal or inland refugia for *Pericopsis elata* and *Erythrophleum suaveolens*. In these species the inland populations that are equally likely to have survived either in the coastal refugia of Cameroon and Gabon or in the inland refugia in Congo (4). Therefore, two additional mixed effects models were run: (iv) LGM-Maley-Coast, including the coastal Maley refugia only, and (v) LGM-Maley-Congo, including the Congo refugia only.

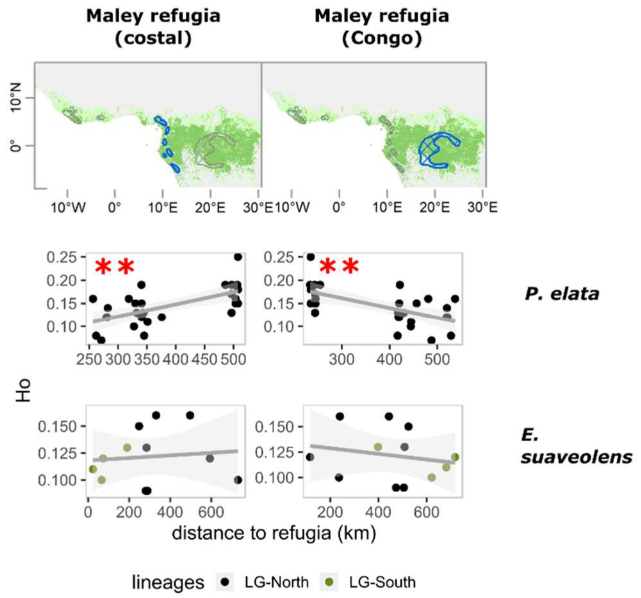

S7. Present-day and Last Glacial Maximum (LGM) niche models for *Pericopsis elata, Distemonanthus benthamianus*, *Erythrophleum ivorense, E. suaveolens*, and *Scorodophloeus zenkeri*. Black dots indicate the present-day ocurrences of individual trees collected from RAINBIO database (5) and our own records (1,029 occurrences in total). Five climatic variables from Worldclim dataset v1.4 at 2.5 arc-min (6) were selected: annual mean temperature (bio1), temperature annual range (bio7), total annual precipitation (bio12), precipitation of the driest month (bio14), and precipitation of December, January and February (pp_djf).

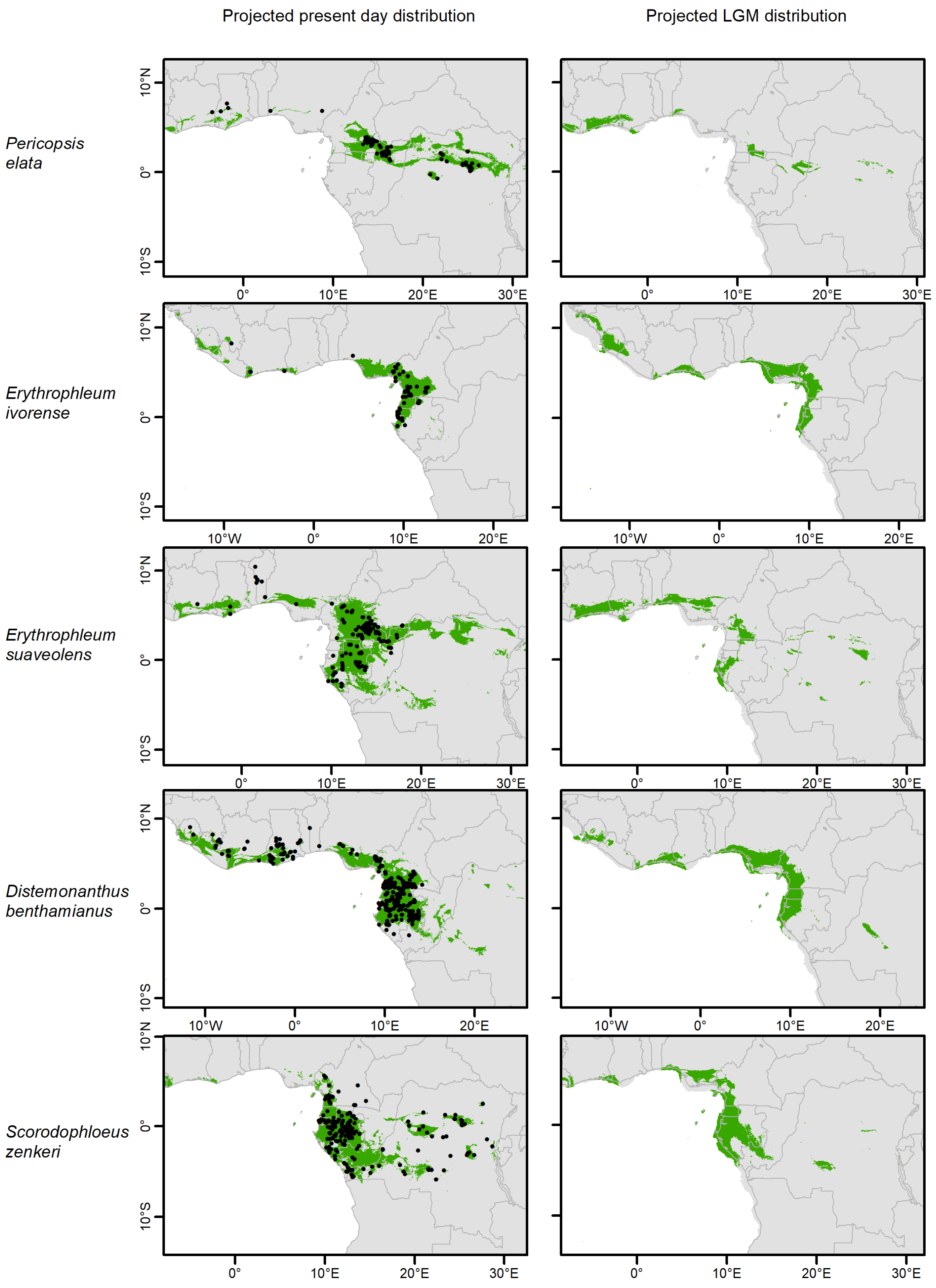

S8. Collection data and genetic cluster based on GBS data of 175 individuals of five Legume tree species widespread in the rainforests of West and Central Africa *Pericopsis elata* (Harms) Meeuwen, *Distemonanthus benthamianus* Baill., *Erythrophleum ivorense* A. Chev., *Erythrophleum suaveolens* (Guill. & Perr.) Brenan,and *Scorodophloeus zenkeri* Harms. Seven outgroup species (OUT) were also sequenced: *Pericopsis laxiflora* (Benth. Ex Baker) Meewen, *Dicorynia guianensis* Amshoff, *Dialium guianense* (Aubl.) Sandwith, *Erythrophleum africanum* (Welw. Ex Benth.) Harms, *Erythrophleum chlorostachys* Baill., *Calpocalyx brevibracteatus* Harms, and *Hymenostegia afzelii* (Oliv.) Harms. The genetic clusters were Upper Guinea (UG), Northern Lower Guinea (nLG), Southern Lower Guinea (sLG), Eastern Lower Guinea (eLG), and Congo.

| SAMPLE_ID | SPECIES | COUNTRY AND LOCALITY | ALTITUDE | LATITUDE | LONGITUDE | LEGIT | DATE | GENETIC CLUSTER |
|---|---|---|---|---|---|---|---|---|
| OH3175 | *P. elata* | Ghana, Afram HdWtrs | | 7.20 | -1.72 | UGaine M | 08/16/2011 | UG |
| NB0570 | *P. elata* | Ghana, Afram Head Waters Forest Reserve | 445 | 5.78 | -0.36 | Bourland N & Dabo J | 05/21/2012 | UG |
| JLD0926 | *P. elata* | Cameroon, Kika | 466 | 2.13 | 15.45 | Ntoude E | 12/28/2009 | nLG |
| JLD0937 | *P. elata* | Cameroon, Socambo | 465 | 2.08 | 15.64 | Ntoude E | 12/28/2009 | nLG |
| JLD0950 | *P. elata* | Cameroon, Salapoumbe | 414 | 2.73 | 15.29 | Ntoude E | 12/28/2009 | nLG |
| CM0050 | *P. elata* | Cameroon, UFA 10030 | | 3.17 | 14.26 | | | nLG |
| NB0224 | *P. elata* | Cameroon, UFA 10030 | | 3.12 | 14.37 | Bourland N | 04/15/2009 | nLG |
| NB0725 | *P. elata* | Cameroon, Pallisco | 700 | 3.94 | 13.83 | Bouissou C | 04/05/2012 | nLG |
| CFC0009 | *P. elata* | Cameroon, Mindourou | | 3.33 | 13.72 | Bourland N | 02/28/2009 | nLG |
| CM0070P | *P. elata* | Cameroon, UFA 10030 | | 3.35 | 14.30 | | | nLG |
| CM0083 | *P. elata* | Cameroon, UFA 10030 | | 3.37 | 14.28 | | | nLG |
| 11/03/2011 | *P. elata* | Cameroon, UFA 10026 Alpicam | 630 | 3.65 | 14.45 | Gillet JF | 07/05/2010 | nLG |
| JLD0035 | *P. elata* | Cameroon, M'bang | | 3.56 | 14.26 | Doucet JL | 02/25/2009 | nLG |
| JLD0928 | *P. elata* | Cameroon, Kika | 378 | 2.09 | 15.54 | Ntoude E | 12/28/2009 | nLG |
| JLD0929 | *P. elata* | Cameroon, Kika | 378 | 2.09 | 15.54 | Ntoude E | 12/28/2009 | nLG |
| JLD0930 | *P. elata* | Cameroon, Kika | 374 | 2.08 | 15.54 | Ntoude E | 12/28/2009 | nLG |
| JLD0936 | *P. elata* | Cameroon, Socambo | 471 | 2.08 | 15.64 | Ntoude E | 12/28/2009 | nLG |
| JLD0938 | *P. elata* | Cameroon, Socambo | 461 | 2.08 | 15.64 | Ntoude E | 12/28/2009 | nLG |
| JLD0942 | *P. elata* | Cameroon, Lobéké | 489 | 2.23 | 15.64 | Ntoude E | 12/28/2009 | nLG |
| JLD0945 | *P. elata* | Cameroon, Lobéké | 487 | 2.23 | 15.64 | Ntoude E | 12/28/2009 | nLG |
| MH1111 | *P. elata* | Cameroon, UFA 30 | | 3.83 | 13.65 | Daïnou K | 08/01/2006 | nLG |
| NB0189 | *P. elata* | Cameroon, UFA 10030 | | 3.12 | 14.33 | Bourland N & al | 04/26/2008 | nLG |
| NB0194 | *P. elata* | Cameroon, UFA 10030 | | 3.14 | 14.33 | Bourland N & al | 04/27/2008 | nLG |

| | | | | | | | | |
|---|---|---|---|---|---|---|---|---|
| NB0198 | *P. elata* | Cameroon, UFA 10030 | | 3.12 | 14.33 | Bourland N & al | 04/27/2008 | nLG |
| NB0370 | *P. elata* | Cameroon, UFA 10038 | 700 | 3.80 | 14.16 | Bourland N | 03/19/2011 | nLG |
| NB0740 | *P. elata* | Cameroon, Pallisco | 700 | 3.91 | 13.82 | Bouissou C | 11/15/2011 | nLG |
| CM0053 | *P. elata* | Cameroon, UFA 10030 | | 3.38 | 13.98 | | | nLG |
| CM0097 | *P. elata* | Cameroon, UFA 10030 | | 3.34 | 14.23 | | | nLG |
| JFG0397 | *P. elata* | Cameroon, UFA 10026 Alpicam | 595 | 3.55 | 14.67 | Gillet JF | 07/03/2010 | nLG |
| JFG0703 | *P. elata* | Cameroon, UFA 10009 SEFAC | 400 | 5.56 | 13.24 | Gillet JF | 06/15/2011 | nLG |
| JLD0932 | *P. elata* | Cameroon, Kika | 422 | 2.04 | 15.62 | Ntoude E | 12/28/2009 | nLG |
| NB0228 | *P. elata* | Cameroon, UFA 10030 | | 3.12 | 14.36 | Bourland N | 04/15/2009 | nLG |
| NB0359 | *P. elata* | Cameroon, UFA 10030 | 655 | 3.89 | 13.60 | Bourland N | 02/21/2011 | nLG |
| NB0392 | *P. elata* | Cameroon, UFA 10031 | | 3.93 | 13.81 | Bourland N | 10/16/2011 | nLG |
| AC0028 | *P. elata* | CongoK, Yoko | 403 | 0.29 | 25.34 | Amani C | 09/01/2010 | CONGO |
| MP0405 | *P. elata* | CongoK, Yangambi | 473 | 0.80 | 24.48 | Pluijgers M | 02/12/2011 | CONGO |
| FBB0171 | *P. elata* | CongoK, BIARO 400 ha | 438 | 0.20 | 25.34 | Boyemba FB | 04/30/2012 | CONGO |
| GiD0508 | *P. elata* | CongoK, Yoko/Kisangani | | 0.30 | 25.30 | Vleminckx J | 08/17/2008 | CONGO |
| NB0100 | *P. elata* | CongoK, Yoko | 450 | 0.29 | 25.32 | Bourland N | 06/05/2009 | CONGO |
| NB0111 | *P. elata* | CongoK, Yoko | 465 | 0.29 | 25.32 | Bourland N | 06/05/2009 | CONGO |
| NB0151 | *P. elata* | CongoK, Yoko | 450 | 0.29 | 25.29 | Bourland N | 06/04/2009 | CONGO |
| NB0289 | *P. elata* | CongoK, Yangambi | 480 | 0.80 | 24.48 | Bourland N | 05/06/2009 | CONGO |
| NB0094 | *P. elata* | CongoK, Yangambi | 465 | 0.81 | 24.48 | Bourland N | 06/07/2009 | CONGO |
| NB0157 | *P. elata* | CongoK, Yangambi | 465 | 0.81 | 24.48 | Bourland N | 06/07/2009 | CONGO |
| MP0359 | *P. elata* | CongoK, Yoko | 436 | 0.30 | 25.29 | Pluijgers M. | 03/11/2011 | CONGO |
| FOLI076 | *P. laxiflora* | Ghana, Kogya nature reserve | 210 | 7.30 | -1.18 | Dabo J | 03/02/2013 | OUT *P. elata* |
| BoD0725 | *D. benthamianus* | Nigeria, Queens Plots Forest Reserved | | 7.19 | 4.97 | Demenou B | 05/03/2014 | DG |
| BoD0296 | *D. benthamianus* | Benin, Forêt de Pobè | | 6.96 | 2.68 | Demenou B | 04/19/2014 | DG |
| BoD0580 | *D. benthamianus* | Ghana, Bowiri | 193 | 7.36 | 0.46 | Demenou B | 04/15/2014 | DG_UG |
| BoD0589 | *D. benthamianus* | Ghana, Nkonia Adenkenso | 136 | 7.26 | 0.33 | Demenou B | 05/15/2014 | DG_UG |
| BoD0525 | *D. benthamianus* | Togo, Badou | 226 | 7.58 | 0.60 | Demenou B | 05/04/2014 | DG_UG |
| MH2688 | *D. benthamianus* | Cameroon, Ngoyang | 512 | 3.34 | 10.74 | Heuertz M | | nLG |
| NB0374 | *D. benthamianus* | Cameroon, UFA 11005 (bordure) | 200 | 5.48 | 8.96 | Bourland N | 07/13/2011 | nLG |
| MH1130 | *D. benthamianus* | Cameroon, UFA 47 | | 3.68 | 13.14 | Daïnou K | 08/01/2006 | nLG |

| | | | | | | | | |
|---|---|---|---|---|---|---|---|---|
| MH2009 | *D. benthamianus* | Cameroon, UFA10 | | 3.82 | 13.33 | NATURE + | | nLG |
| OH1929 | *D. benthamianus* | Cameroon, route Eboumetoum | 704 | 3.68 | 13.14 | Emerano Z | 08/14/2007 | nLG |
| NB0421 | *D. benthamianus* | Cameroon, Djoum | | 2.70 | 12.65 | Bourland N | 11/21/2011 | nLG |
| OH3355 | *D. benthamianus* | Cameroon | 684 | 2.64 | 12.66 | Hardy O | 10/27/2013 | nLG |
| OH3358 | *D. benthamianus* | Cameroon | 697 | 2.51 | 12.67 | Hardy O | 10/27/2013 | nLG |
| JLD1055 | *D. benthamianus* | Cameroon, Sangmelima | 675 | 3.06 | 12.03 | Ntoude E | 01/11/2010 | nLG |
| MH1450 | *D. benthamianus* | Cameroon, Djoum-Oveng-Sangmélima | 620 | 2.38 | 13.43 | Heuertz M | 07/11/2007 | nLG |
| JLD1032 | *D. benthamianus* | Cameroon, Djoum | 638 | 2.69 | 12.91 | Ntoude E | 01/09/2010 | nLG |
| MH1468 | *D. benthamianus* | Cameroon, Djoum-Oveng-Sangmélima | 624 | 2.60 | 12.10 | Heuertz M | 07/11/2007 | nLG |
| JLD1057 | *D. benthamianus* | Cameroon, Zoétélé | 658 | 3.31 | 11.80 | Ntoude E | 01/11/2010 | nLG |
| NB0327 | *D. benthamianus* | Cameroon, UFA 09-021 | | 2.45 | 10.64 | Bourland N | 06/17/2010 | nLG_sLG |
| MH1575 | *D. benthamianus* | Cameroon, route Okong-Messama-Mefo | 582 | 2.59 | 10.90 | Heuertz M | 07/22/2007 | nLG_sLG |
| OH1876 | *D. benthamianus* | Gabon, Ebe-Messe | 483 | 0.29 | 12.09 | Koumba P | 02/26/2008 | sLG |
| OH3064 | *D. benthamianus* | Gabon, Sud Mt Cristal (Rougier-Ht Abanga) | 256 | 0.29 | 11.21 | Hardy O | 05/19/2011 | sLG |
| OH3072 | *D. benthamianus* | Gabon, Sud Mt Cristal (Rougier-Ht Abanga) | 325 | 0.16 | 11.13 | Hardy O | 05/19/2011 | sLG |
| OH1804 | *D. benthamianus* | Gabon, Okondja | | -0.95 | 13.63 | Boubady AG | 10/10/2007 | sLG |
| OH1435 | *D. benthamianus* | Gabon, CFAD CEB | | -0.79 | 12.85 | Boubady AG | | sLG |
| OH1701 | *D. benthamianus* | Gabon, route Bambidie-Lelama | 290 | -0.94 | 13.37 | Boubady AG | 02/18/2008 | sLG |
| OH1437 | *D. benthamianus* | Gabon, CFAD CEB | | -0.55 | 12.83 | Boubady AG | | sLG |
| OH2683 | *D. benthamianus* | Gabon, Mt de Cristal | 471 | 1.00 | 10.92 | Hardy O | 05/12/2011 | sLG |
| OH1449 | *D. benthamianus* | Gabon, Nonié | 33 | -0.02 | 9.37 | Doucet JL | | sLG_nLG |
| OH1450 | *D. benthamianus* | Gabon, Nonié | 33 | -0.02 | 9.37 | Doucet JL | | sLG_nLG |
| OH1483 | *D. benthamianus* | Gabon, CBG (côte sud) | 56 | -1.70 | 10.17 | Laporte J | | sLG_nLG |
| OH1464 | *D. benthamianus* | Gabon, CBG (côte sud) | 144 | -1.83 | 10.46 | Laporte J | | sLG_nLG |
| GK1030 | *D. benthamianus* | Ivory Coast, Scio | 436 | 6.42 | -7.48 | Koffi G | 01/27/2009 | UG |
| GK1031 | *D. benthamianus* | Ivory Coast, Scio | 470 | 6.42 | -7.48 | Koffi G | 01/27/2009 | UG |
| OH3220 | *D. benthamianus* | Ghana, Nkrabia Road | 132 | 6.01 | -1.55 | Hardy OJ | 03/26/2013 | UG |
| OH3222 | *D. benthamianus* | Ghana, Oda-Edubiase road | 141 | 6.03 | -1.33 | Hardy OJ | 03/27/2013 | UG |
| OH3231 | *D. benthamianus* | Ghana, Road Techiman-Sunyani | 340 | 7.50 | -2.09 | Hardy OJ | 03/05/2013 | UG |
| BoD0658 | *D. benthamianus* | Ghana, Yoyo Forest River | 135 | 5.92 | -2.83 | Demenou B | 05/19/2014 | UG |
| BoD0638 | *D. benthamianus* | Ghana, Ankasa Forest | 34 | 5.28 | -2.74 | Demenou B | 05/18/2014 | UG |

| | | | | | | | | |
|---|---|---|---|---|---|---|---|---|
| MP0011 | *D. benthamianus* | Ghana, Boabeng | | 7.72 | -1.69 | | 07/13/2010 | UG |
| MP0043 | *D. benthamianus* | Ghana, Bunso (arboretum domain) | | 5.76 | -0.22 | | 07/30/2010 | UG |
| NB0578 | *D. benthamianus* | Ghana | 325 | 7.09 | -1.72 | Bourland N | 05/21/2012 | UG |
| LV1094414 | *D. guianensis* | Guyane, Saut Lavilette_SE of Regina | 187 | -52.20 | 4.15 | C Baraloto, J Chave & al | 09/01/2008 | OUT *D. benthamianus* |
| EE0426 | *D. guianense* | Bénin, Sarimanga / Nagayilè - Donga | | 9.25 | 1.83 | Ewédjè EE | 05/03/2009 | OUT *D. benthamianus* |
| JD0065 | *E. ivorense* | Cameroon, Korup plot | | 8.83 | 5.06 | Duminil J | | nLG |
| OH0693 | *E. ivorense* | Cameroon, Korup plot | | 8.85 | 5.07 | Hardy OJ | 03/10/2006 | nLG |
| OH2029 | *E. ivorense* | Cameroon | 155 | 9.07 | 5.61 | Hardy OJ | 04/25/1901 | nLG |
| OH0518 | *E. ivorense* | Cameroon, Limbe, Bimbia Community Forest | | 9.10 | 4.01 | Hardy OJ | 03/03/2006 | nLG |
| TOD1746 | *E. ivorense* | Cameroon, UFA 11-001 (TRC) | | 9.22 | 5.46 | Doumenge C | 11/30/1910 | nLG |
| MH1809 | *E. ivorense* | Cameroon, Ma'an | | 10.62 | 2.39 | Daïnou K | 07/22/2007 | nLG_sLG |
| JD0053 | *E. ivorense* | Cameroon, Korup plot | | 8.84 | 5.06 | Duminil J | | nLG |
| SVO0073 | *E. ivorense* | Cameroon, Mekas, vers la rivière Ongwene | 635 | 12.54 | 3.16 | Onana J-M | 03/09/1901 | nLG |
| NB0238 | *E. ivorense* | Cameroon, UFA 09-021 | | 10.65 | 2.46 | Bourland N | 05/08/2009 | nLG_sLG |
| OH0716 | *E. ivorense* | Cameroon, Korup NP | | 8.84 | 5.06 | Hardy OJ | | nLG |
| GID0850 | *E. ivorense* | Gabon, Forêt classée de la Mondah | | 9.33 | 0.58 | Dauby G | 02/28/2007 | sLG |
| CD0363 | *E. ivorense* | Gabon | | 9.58 | 0.96 | Doumenge C | | sLG |
| CD0427 | *E. ivorense* | Gabon, Mvam, CFAD de Rimbunan Ijau | | 9.65 | -0.18 | Doumenge C | 06/20/2008 | sLG |
| MH1811 | *E. ivorense* | Cameroon, Ile Ipika | 62 | 9.95 | 2.28 | Daïnou K | 07/25/2007 | nLG_sLG |
| MH0722 | *E. ivorense* | Gabon, Pointe Denis | | 9.35 | 0.34 | Heuertz M | 08/27/2006 | sLG |
| GK0662 | *E. ivorense* | Ivory Coast, Haute Dodo | 436 | -7.05 | 5.05 | Koffi G | | UG |
| LK0017 | *E. ivorense* | Ivory Coast, Mélékoukro | 5 | -3.29 | 5.16 | Kouadio L | 11/25/1910 | UG |
| EE0662 | *E. ivorense* | Nigeria, Ilesha | 383 | 6.09 | 6.22 | Ewédjè EE | 08/16/1911 | nLG |
| JD0507 | *E. suaveolens* | Cameroon, Forêt communale de Gari | | 14.17 | 4.71 | Duminil J | 02/15/2007 | nLG |
| JD0584 | *E. suaveolens* | Cameroon, UFA10-031 | | 14.27 | 3.37 | NATURE + | | nLG |
| JD0713 | *E. suaveolens* | Cameroon, UFA 10 041 | | 13.80 | 3.21 | NATURE + | 10/16/1910 | nLG |
| OH1387 | *E. suaveolens* | Cameroon, Forêt communautaire de Kongoulou | | 13.82 | 3.02 | Doumenge C | 03/07/2007 | nLG |
| JD0585 | *E. suaveolens* | Cameroon, UFA10-031 | | 13.86 | 3.80 | NATURE + | | nLG |
| JD0566 | *E. suaveolens* | Cameroon, UFA31 | | 13.90 | 3.80 | Duminil J | 02/16/2007 | nLG |
| JFG0024 | *E. suaveolens* | CongoB, CIB Pokola | 34 | 16.40 | 1.32 | Gillet JF | 07/05/2008 | nLG |

| | | | | | | | | |
|---|---|---|---|---|---|---|---|---|
| JD0676 | *E. suaveolens* | Cameroon, UFA 10 044 | 71 | 13.52 | 3.62 | NATURE + | 10/08/1910 | nLG |
| OH1414 | *E. suaveolens* | Cameroon, UFA 10-063 d'Alpicam | | 15.53 | 2.02 | Doumenge C | 03/12/2007 | nLG |
| JD0660 | *E. suaveolens* | CongoB, Pokola | 33 | 16.39 | 1.32 | Gillet JF | 10/27/1910 | nLG |
| KD0298 | *E. suaveolens* | Central African Republic, Lolé | 531 | 17.87 | 3.83 | Kasso D | | nLG |
| RM0010 | *E. suaveolens* | Gabon, SEEF/Milolé | | 13.14 | -0.29 | Mboma R | 05/26/2009 | sLG |
| CD0152 | *E. suaveolens* | Gabon | | 10.16 | -0.90 | Doumenge C | | sLG |
| CD0478 | *E. suaveolens* | Gabon, Tchibanga-Ndendé, Mouindji, Nyanga | | 11.13 | -2.72 | Doumenge C | 06/23/2008 | sLG |
| CD0328 | *E. suaveolens* | Gabon | | 10.46 | -1.46 | Doumenge C | | sLG |
| TOD0931 | *E. suaveolens* | Cameroon, Sommet falaise Djandjou | | 11.74 | 5.43 | Arbonnier M | 07/28/2008 | UG |
| LK0033 | *E. suaveolens* | Ivory Coast, Ahrémou II | 95 | -4.93 | 6.21 | Kouadio L | 01/10/2009 | UG |
| LK0040 | *E. suaveolens* | Ivory Coast, Ahrémou II | 86 | -4.92 | 6.22 | Kouadio L | 01/10/2009 | UG |
| MH2260 | *E. suaveolens* | Benin, Assanhoun | | 1.66 | 8.62 | González-Martínez S | | UG |
| EE0282 | *E. suaveolens* | Benin, Forêt classée de Soudou,anigri | 327 | 1.77 | 8.90 | Ewédjè EE | 11/30/1910 | UG |
| EE0463 | *E. suaveolens* | Benin, Igbo Aladja | 252 | 2.23 | 8.75 | Ewédjè EE | 04/24/2009 | UG |
| MH2279 | *E. suaveolens* | Benin, Etchede | | 2.63 | 6.99 | González-Martínez S | | UG |
| TOD0786 | *E. suaveolens* | Cameroon, Mankumbe | | 11.13 | 5.85 | Arbonnier M | 07/15/2008 | UG |
| OH1384 | *E. suaveolens* | Cameroon, Savanes au Nord de la Sanaga, Yaoundé-Bafia | | 11.26 | 4.48 | Doumenge C | 03/04/2007 | UG |
| EE299 | *E. africanum* | Benin, Forêt galerie nettement isolée | 513 | 10.21 | 1.21 | Ewédjè EE | 14/03/2009 | OUT *Erythrophleum* sp. |
| EC | *E. chlorostachys* | Australia | | -12.49 | 130.99 | | | OUT *Erythrophleum* sp. |
| WHA0095 | *C. brevibracteatus* | Ghana | 118 | 5.59 | -2.43 | Hawthorne WD | 15/07/2013 | OUT *Erythrophleum* sp. |
| NB0148 | *S. zenkeri* | CongoK, Yoko | 445 | 0.29 | 25.30 | Bourland N | 06/04/2009 | CONGO |
| NB0172 | *S. zenkeri* | CongoK, Yangambi | 475 | 0.81 | 24.49 | Bourland N | 06/07/2009 | CONGO |
| NB0281 | *S. zenkeri* | CongoK, Yoko | 450 | 0.29 | 25.32 | Bourland N | 06/05/2009 | CONGO |
| NB0292 | *S. zenkeri* | CongoK, Yangambi | 470 | 0.80 | 24.49 | Bourland N | 05/06/2009 | CONGO |
| AD0700 | *S. zenkeri* | Gabon, Makokou, For t dense sempervirente | 678 | 0.91 | 13.67 | Donkpegan A | 05/20/2014 | eLG |
| GiD0605 | *S. zenkeri* | Gabon, Concession Rougier de l'Ivindo, | 311 | -0.08 | 12.37 | Dauby G | 03/19/2009 | eLG |
| GiD0799 | *S. zenkeri* | Gabon, Concession Rougier de l'Ivindo | 309 | -0.08 | 12.35 | Dauby G | 03/17/2009 | eLG |
| GiD0968 | *S. zenkeri* | Parc National de Minkébé, amont de la rivière Sing | 500 | 1.50 | 12.81 | Dauby G | 08/11/2009 | eLG |
| GiD1041 | *S. zenkeri* | Parc National de Minkébé, amont de la rivière Sing | 484 | 1.13 | 13.02 | Dauby G | 08/09/2009 | eLG |
| GiD1224 | *S. zenkeri* | Gabon, Sud-Est de Koulamoutou | 562 | -1.34 | 12.71 | Dauby G | 02/22/2009 | eLG |

| | | | | | | | | |
|---|---|---|---|---|---|---|---|---|
| GK0121 | *S. zenkeri* | Gabon, Makokou - IRET | 502 | 0.52 | 12.79 | Koffi G | 11/06/2007 | eLG |
| GK0363 | *S. zenkeri* | Gabon, Kongou | 501 | 0.29 | 12.57 | Koffi G | 11/26/2007 | eLG |
| OH0114 | *S. zenkeri* | Gabon, Station IRET- Makokou | 524 | 0.01 | 12.80 | Hardy O | 08/04/2005 | eLG |
| RM0031 | *S. zenkeri* | Gabon, Mitzic(Foreex/Madouaka) | | 0.57 | 12.05 | Mboma R | 03/24/2009 | eLG |
| RM0036 | *S. zenkeri* | Gabon, Mitzic(Foreex/Madouaka) | | 0.58 | 12.07 | Mboma R | 03/25/2009 | eLG |
| RM0039 | *S. zenkeri* | Gabon, SEEF/Milolé | | -0.42 | 12.85 | Mboma R | 06/01/2009 | eLG |
| RM0044 | *S. zenkeri* | Gabon, SEEF/Milolé | | -0.41 | 12.87 | Mboma R | 06/02/2009 | eLG |
| RM0047 | *S. zenkeri* | Gabon, SEEF/Milolé | | -0.30 | 13.11 | Mboma R | 06/03/2009 | eLG |
| MH1627 | *S. zenkeri* | Cameroon, Nkong Mekak | 478 | 2.79 | 10.54 | | 07/22/2007 | nLG |
| MH1631 | *S. zenkeri* | Cameroon, Nkong Mekak | 489 | 2.79 | 10.54 | | 07/23/2007 | nLG |
| MH1912 | *S. zenkeri* | Cameroon, Akom II | 753 | 3.26 | 10.91 | Nguembou C | 06/25/2007 | nLG |
| MH1921 | *S. zenkeri* | Cameroon, Bifa | 82 | 3.11 | 10.47 | Nguembou C | 06/13/2007 | nLG |
| MH2667 | *S. zenkeri* | Cameroon, Ngoyang | 488 | 3.35 | 10.75 | Heuertz H, Budde K, Mambo | | nLG |
| MH2678 | *S. zenkeri* | Cameroon, Ngoyang | 465 | 3.34 | 10.74 | Heuertz H, Budde K, Mambo | | nLG |
| OH0883 | *S. zenkeri* | Cameroon, Akom | 640 | 2.75 | 10.54 | Droissart V | 06/05/2006 | nLG |
| TOD0567 | *S. zenkeri* | Cameroon, UFA 00-003 (MMG) | | 3.39 | 10.43 | | 02/14/2008 | nLG |
| TOD0665 | *S. zenkeri* | Cameroon, Massif de Ngovayang, Mvilé | | 3.24 | 10.58 | | 02/18/2008 | nLG |
| GiD0472 | *S. zenkeri* | Gabon, Zone d'exploration de Sogademin | 50 | 0.45 | 10.18 | Dauby G | 09/03/2008 | sLG1 |
| GiD0604 | *S. zenkeri* | Gabon, Nord de Ndjolé, concession de l'Ogooué | 194 | -0.09 | 10.76 | Dauby G | 02/22/2009 | sLG1 |
| GiD1002 | *S. zenkeri* | Gabon, Zone d'exploration de Sogademin | 88 | 0.46 | 10.19 | Dauby G | 07/14/2009 | sLG1 |
| OH3111 | *S. zenkeri* | Gabon, Sud Mt Cristal (Rougier-Ht Abanga) | 117 | -0.16 | 10.63 | Hardy O | 05/19/2011 | sLG1 |
| TS0106 | *S. zenkeri* | Gabon, Concession SEEF, Monts de Cristal | 735 | 0.48 | 10.50 | Stevart T | 10/24/2010 | sLG1 |
| GiD0791 | *S. zenkeri* | Gabon, Concession Rougier du Haut-Abanga | | 0.43 | 11.21 | Nguema D | 11/21/2008 | sLG1_eLG |
| OH2879 | *S. zenkeri* | Gabon, Woleu Ntem | 346 | 0.53 | 11.47 | Hardy O | 05/16/2011 | sLG1_eLG |
| OH3092 | *S. zenkeri* | Gabon, Sud Mt Cristal (Rougier-Ht Abanga) | 121 | -0.06 | 10.98 | Hardy O | 05/19/2011 | sLG1_eLG |
| GiD0612 | *S. zenkeri* | Gabon, CFAD de Rimbunan Hijau | 423 | -0.87 | 11.26 | Dauby G | 02/02/2009 | sLG1_sLG2 |
| GiD0797 | *S. zenkeri* | Gabon, CFAD de Rimbunan Hijau | 373 | -0.85 | 11.27 | Dauby G | 01/27/2009 | sLG1_sLG2 |
| GiD1715 | *S. zenkeri* | Gabon, Zone de Mabounié, ca. 45 km de Lambaréné | 100 | -0.77 | 10.54 | Dauby G | 05/07/2012 | sLG1_sLG2 |
| AD0740 | *S. zenkeri* | Gabon, Mayumba | 263 | -3.28 | 10.84 | Donkpegan A | 06/03/2014 | sLG2 |
| GiD0895 | *S. zenkeri* | Gabon, Tchibanga | 475 | -3.28 | 11.12 | Nguema D | 04/06/2009 | sLG2 |

| | | | | | | | | |
|---|---|---|---|---|---|---|---|---|
| GiD1105 | *S. zenkeri* | Gabon, Waka river | 163 | -1.39 | 10.86 | Stevart T | 11/11/2009 | sLG2 |
| MH0996 | *S. zenkeri* | Gabon, Douano | 386 | -2.64 | 11.25 | Heuertz M | 09/13/2006 | sLG2 |
| TS0147 | *S. zenkeri* | Gabon, Mt Birougou | 780 | -2.04 | 12.23 | Stevart T | 02/15/2011 | sLG2 |
| MH0743 | *S. zenkeri* | Gabon, Popa, pr. Michimba et du PN Mont Birougou | 714 | -1.61 | 12.27 | Heuertz M | 09/01/2006 | sLG2_eLG |
| MH0753 | *S. zenkeri* | Gabon, Popa, pr. Michimba et du PN Mont Birougou | 675 | -1.62 | 12.28 | Heuertz M | 09/01/2006 | sLG2_eLG |
| GK0193 | *S. zenkeri* | Gabon, Bélinga, forêt mature; terre ferme; pente forte | 640 | 1.11622 | 13.22166 | Koffi G | 16/11/2007 | eLG |
| MH0971 | *Crudia gabonensis* | Gabon, Douano, Forêt primaire sur sol bien drainé | 374 | -2.6756 | 11.2509 | Heuertz M | 12/09/2006 | OUT *S. zenkeri* |

### SI REFERENCES


1. Hawthorne W (1995) Ecological profiles of Ghanaian forest trees. *Tropical forestry papers* (29).
2. Hardy OJ*, et al.* (2013) Comparative phylogeography of African rain forest trees: a review of genetic signatures of vegetation history in the Guineo-Congolian region. *Comptes Rendus Geoscience* 345(7-8):284-296.
3. Hardy OJ*, et al.* (2019) Seed and pollen dispersal distances in two African legume timber trees and their reproductive potential under selective logging. *Molecular ecology* 28:3119-3134.
4. Maley J (1996) The African rain forest–main characteristics of changes in vegetation and climate from the Upper Cretaceous to the Quaternary. *Proceedings of the Royal Society of Edinburgh, Section B: Biological Sciences* 104:31-73.
5. Dauby G*, et al.* (2016) RAINBIO: a mega-database of tropical African vascular plants distributions. *PhytoKeys* (74):1.
6. Hijmans RJ, Cameron SE, Parra JL, Jones PG, & Jarvis A (2005) Very high resolution interpolated climate surfaces for global land areas. *International journal of climatology* 25(15):1965-1978.
